# A pro-endocrine pancreatic transcriptional program established during development is retained in human gallbladder epithelial cells

**DOI:** 10.1101/2021.03.02.433636

**Authors:** Mugdha V. Joglekar, Subhshri Sahu, Wilson KM Wong, Sarang N. Satoor, Charlotte X. Dong, Ryan J Farr, Michael D. Williams, Prapti Pandya, Gaurang Jhala, Sundy N.Y. Yang, Yi Vee Chew, Nicola Hetherington, Dhan Thiruchevlam, Sasikala Mitnala, Guduru V Rao, Duvvuru Nageshwar Reddy, Thomas Loudovaris, Wayne J. Hawthorne, Andrew G. Elefanty, Vinay M. Joglekar, Edouard G. Stanley, David Martin, Helen E. Thomas, David Tosh, Louise T. Dalgaard, Anandwardhan A. Hardikar

## Abstract

**Objective:** Pancreatic islet β-cells are factories for insulin production; however ectopic expression of insulin is also well recognized. The gallbladder is a next-door neighbour to the developing pancreas. Here, we wanted to understand if gallbladders contain functional insulin-producing cells.

**Design:** We compared developing and adult mouse as well as human gallbladder epithelial cells and islets using immunohistochemistry, flow cytometry, ELISAs, RNA-sequencing, real-time PCR, chromatin immunoprecipitation and functional studies.

**Results:** We demonstrate that the epithelial lining of developing, as well as adult mouse and human gallbladders naturally contain interspersed cells that retain the capacity to actively transcribe, translate, package, and release insulin. We show for the first time that human gallbladders also contain functional insulin-secreting cells with the potential to naturally respond to glucose *in vitro* and *in situ*. Notably, in a NOD mouse model of type 1 diabetes, we observed that insulin-producing cells in the gallbladder are not targeted by autoimmune cells. Conclusion: In summary, our biochemical, transcriptomic, and functional data in human gallbladder epithelial cells collectively demonstrate their potential for insulin-production under pathophysiological conditions, and open newer areas for type 1 diabetes research and therapy.

**Significance of the study:** 

**What is already known about this subject?:**

- Developing pancreas and gallbladder are next-door neighbours and share similar developmental pathways.
- Human Gallbladder-derived progenitor cells were shown to differentiate into insulin-producing cells.

**What are the new findings?:**

- Gallbladder epithelium contains interspersed cells that can transcribe, translate, package and secrete insulin.
- Insulin-producing cells in the gallbladder are not destroyed by immune cells in an animal model of type 1 diabetes (T1D).
- Our studies demonstrating the absence of insulin splice variants in human gallbladder cells, and higher splice forms in human islets, suggest a potential mechanism (via defective ribosomal products) in escaping islet autoimmunity.

**How might it impact clinical practice?:**

- Deciphering mechanisms of protection of insulin-producing cells from immune cells in the gallbladder could help in developing strategies to prevent islet autoimmunity in T1D.

## Introduction

The regulation of mammalian gene expression is an important and complex issue in biology. Although pancreatic islet β-cells are the major insulin-producing cells in our body, we and others have demonstrated ectopic expression of insulin in the gallbladder/biliary duct epithelium ^1 2^, thymus ^3^, and the brain ^4 5^. The gallbladder and pancreas arise from the foregut endoderm and share transcription factors during embryonic development ^6^. The occurrence of β-like cells in mouse biliary ducts was reported earlier ^7^ and the inhibition of *Hes1* in mouse gallbladder epithelial cells improved insulin production ^2^. A recent study also implicated therapeutic potential by forced expression of pancreatic transcription factors to differentiate gallbladder-derived cells from individuals with type 1 diabetes (T1D) into insulin-producing cells ^8^.

Here, using mouse models and human tissue/samples spanning embryonic to post-natal/adult life, we demonstrate for the first time, a comparative molecular and functional analysis of insulin-transcribing cells in the gallbladder and pancreas. Whilst expression of genes and their splice variants across human tissues is fundamental to the understanding of tissue specificity and function, current tissue expression datasets, such as the GTEx ^9^, lack gene expression data from the gallbladder. Our work adds new knowledge while demonstrating the gallbladder to be an important and interesting naturally occurring source of insulin-producing cells.

## RESEARCH DESIGN AND METHODS

### Animals

All animal experimentation was carried according to guidelines outlined by the respective institute’s animal care and use committees in India, UK and Australia. Developing pancreatic buds/pancreas and gallbladder tissues were dissected without cross-contamination under a fluorescent-stereomicroscope and samples collected in Trizol or 4% freshly prepared paraformaldehyde.

### Human tissue collection

Gallbladder, pancreas and human islet samples were stored for RNA isolation (in Trizol), fixed (in 4% paraformaldehyde) for immunostaining or processed immediately for cell culture as per human research ethics committee (HREC) approvals. *In situ* glucose stimulation in consented five individuals undergoing cholecystectomies at the Asian Institute of Gastroenterology was performed according to the Declaration of Helsinki II with participants providing informed written consent as per the ethical committee approval from the Asian Institute of Gastroenterology, India.

### Gallbladder cell culture

Gallbladder tissues collected in M199 media were washed thoroughly to remove bile and the inner surface was gently scraped with a disposable scalpel to isolate epithelial lining. Cells were plated in a serum-containing medium (10% FBS + M199 with 25mM HEPES and 2mM glutamine + Hams F12k) with penicillin, streptomycin (Gibco, Carlsbad, CA). This medium is hereafter referred to as a growth-promoting medium or serum-containing medium (SCM).

### *In vitro* differentiation

Human adult gallbladder-derived mesenchymal-like cells or transduced cells, were differentiated as described earlier ^10 11^ with islet-like cell aggregates (ICAs) harvested for RNA isolation by day(D)14. In some experiments, DNMT/HDAC inhibitors were added during differentiation; 1 mM Sodium butyrate (SB), 100nM Trichostatin A (TSA), 1mM Valproic acid (VPA), 2μM 5-aza-2’-deoxycytidine (5-Aza) and 1μM dexamethasone (dex) with appropriate vehicle control (PBS/DMSO). Adenoviral vectors for *PDX1*, *NEUROG3,* and *MAFA* were used at two different multiplicity of infection (MOI). Transductions were performed as described earlier ^12^.

### RNA isolation, cDNA synthesis, and quantitative real-time PCR

RNA isolation was carried using Trizol (Invitrogen, Carlsbad, CA) and quantitative real-time PCR was performed with TaqMan primer-probe mix (**Table S1**) from Thermo Fisher Scientific, Foster City, CA. The cycle threshold (Ct) values were normalized to the housekeeping gene 18S rRNA. Transcript abundance was calculated as described earlier, with Ct-value=39 as the limit of detection for TaqMan-qPCR platform used^13^. TaqMan Low-Density Array (TLDA) cards were used for selected gene panels (**Table S2**) on the 7900 HT system (Thermo Fisher Scientific, Foster City, CA). Normalized Ct-values were used to plot hierarchical cluster heatmaps. Customized OpenArray™ Human mRNA panel (Thermo Fisher Scientific, Waltham, MA) analysed 45 selected pancreatic gene transcripts (**Table S3**). Custom panels were used as per the manufacturer’s protocol on QuantStudioTM 12K Flex Real-Time PCR platform (Thermo Fisher Scientific, Waltham, MA).

### Bulk RNA sequencing

RNA sequencing on mouse tissues was performed using Ion Torrent PGM Instrument (Thermo Fisher Scientific, Waltham, MA) as detailed in^14^. Adult human pancreatic islets and gallbladder epithelial cell samples (gallbladder dataset GSE152419; n=7 and human islet dataset GSE152111, n=66) were sequenced on the HiSeq4000 platform (detailed in ^15^ and SOM). Strand Next-generation sequencing (NGS) v2.5 software was used to analyze the RNA-Seq data as detailed in SOM.

### Pancreatic single-cell (sc)RNA sequencing analysis

Panc8 single-cell sequencing data (n=14,890) was extracted from available datasets (GSE84133, GSE85241, E-MTAB-5061, GSE83139, GSE81608) and analyzed via R studio version 1.2.5033 (built under the R 3.5.1) using the SeuratData (version 0.2.1), Seurat, ggplot2 and cowplot packages as detailed in SOM.

### Immunostaining

Tissue sections/cells were immunostained as described in ^16^ and SOM. Images were scanned and assessed using a Zeiss LSM 510 laser confocal microscope with identical laser/PMTsettings (across samples) and thresholds below saturation.

### Immuno-electron microscopy

Insulin secretory granules were visualized in CD1 mouse gallbladder/pancreas from the same animals using transmission electron microscopy (Jeol 1200EX TEM, University of Bath, UK) as detailed in SOM. Briefly, fixed tissues were blocked in 10% normal goat serum, immunostained using guinea pig anti-Insulin (Linco Research Inc, St. Charles, MO, 1:500 dilution) overnight at 4°C and then gold tagged goat anti-guinea pig antibody (British Biolcell, Cardiff, UK) at 1:50 dilution for 2 hours at room temperature before imaging.

### Flow cytometry

Cells were visualized using a FACSCalibur™ system. Phycoerythrin (PE) labelled Rat IgG2 (isotype control), anti-CD29, anti-CD44, anti-CD90, and anti-CD105 (all PE-labelled BD Biosciences, Franklin Lakes, NJ) at dilutions recommended by the manufacturer. Data were analyzed using CellQuest-Pro software after excluding PI^+^ (dead) cells. Isotype control or wild type tissues were used to set up gating with identical acquisition parameters.

### Cell lineage tracing

*In vitro* propagation of insulin-positive cells was carried out using two thymidine analogues CldU and IdU (Sigma-Aldrich, St Louis, MO) as described before^17^ and detailed in SOM.

### Chromatin immunoprecipitation (ChIP)

Adult mouse gallbladder and pancreas, freshly isolated epithelial cells from the adult human gallbladder, freshly isolated human islets, and gallbladder-derived mesenchymal cells in culture at passage 5 were used in ChIP assay as described before^16^. Immunoprecipitation was performed using 2μg of specific dimethyl-/trimethyl-antibodies for H3K4 and H3K9; acetylation of H3K9 and H3 and H4 (Millipore, Billerica, MA). Input, immunoprecipitated and isotype control DNA was resuspended in nuclease-free water and used for SYBR green or TaqMan qPCR (with appropriate FAST SYBR/TaqMan mastermix) using primers listed in **Table S4** on a ViiA7™ Real-Time PCR System 96-well platform (Thermo Fisher Scientific, Waltham, MA).

### Glucose stimulated insulin secretion

Gallbladder epithelial clusters were handpicked under a phase-contrast microscope, exposed to basal (2.5mM Glucose) or stimulated (25mM Glucose) in KRBH buffer, for 1 hour at 37°C. The supernatant, or cell pellets (obtained by sonicating in 200-500μL of acid ethanol), were measured for insulin/C-peptide content by ELISA kit (Mercodia, Winston Salem, NC).

### Transplantation of human gallbladder epithelial cells

Transplantation of freshly isolated gallbladder epithelial cells was carried out on 8-12 week-old male NOD/SCID to assess their function in response to glucose stimulation as described ^18 19^ and detailed in SOM.

### Pathway analysis

Gallbladder expressed genes (GSE152419) were compared with β-cell-expressed genes (from E-GEOD-20966) using GO-analysis on Pantherdb.org^20^ as detailed in SOM. Gene lists of 16,584 gallbladder transcripts and 13,164 β-cell transcripts were compared for overlaps in transcripts using Venn diagrams (https://bioinfogp.cnb.csic.es/tools/venny/index.html), identifying 7,227 genes to be common. Statistical overrepresentation was calculated using Fisher’s exact test, using Bonferroni correction for multiple testing.

### Statistical analysis

Statistical analyses were performed using GraphPad Prism 8.4.1, the R-software (ver. 3.6.2), SPSS Statistics 27 or Microsoft Excel (ver. 2016). R-software was used to perform unsupervised hierarchical clustering maps using heatmap.2 function in gplots. Add-on and other R-packages XLconnect (with ActivePerl software) and RColorBrewer were used for dataset import and visualization along with R-package gplots. Kolmogorov-Smirnov test was used to check for data normality in SPSS. F-test was performed to check for variance in Excel. For non-normally distributed data, two-tailed Mann-Whitney test was used to calculate the P-value with no ties computed as performed in R. Normally distributed data with equal or unequal variance were analyzed using two-tailed Student’s or Welch’s t-test to calculate the P-value in Excel, respectively. Volcano plots were generated in Excel. GraphPad Prism was used to perform all remaining analyses using appropriate statistical tests and corrected for multiple comparisons if required. Split volcano and spider/radar plots were created using BioVinci data visualization package.

## Results

### Insulin expression in developing mouse gallbladder cells

The gallbladder originates from the pancreatic bud that also gives rise to the ventral pancreas ^6^ (**Fig. 1A**) during embryonic development. We profiled key endocrine pancreatic gene transcripts in developing mouse pancreas and gallbladder at embryonic days(E) 15.5, E16.5, and E18.5. The developing pancreas contained more copies of *Ins1*, *Ins2* (**Fig. 1B**) and *Gcg, Sst* (**Fig. S1A**) gene transcripts as compared to those in the gallbladder. Interestingly, the master regulatory pancreatic transcription factor *Pdx1* was expressed in gallbladder cells at all times, and at levels similar to those in the pancreas closer to birth (E16.5 and E18.5; **Fig. 1B**). The expression of other gene transcripts (*Hnf1b, Sox9, Hnf4a, Hlx*) did not exhibit significant differences between developing pancreas and gallbladder (**Fig. S1A**) during the time points assessed. Developing mouse gallbladders also contain immune-reactive insulin protein, albeit at significantly lower levels than those in the developing pancreas (**Fig. S1B**). Using a *Pdx1*^GFP/w^ reporter mouse^21^ **(Fig. 1C)** we confirmed that cells within the prospective gallbladder and pancreatic buds exhibit *Pdx1* gene promoter activity **(Fig. 1D)**. As expected, wild type (WT) gallbladder cells do not contain any GFP^+^ cells (right bottom; **Fig. 1D).** Chromatin immunoprecipitation (ChIP)^22^ analysis at the *Pdx1* gene promoter region demonstrated that all three sites assessed are active in the gallbladder (**Fig. S1C**). Bulk RNA sequencing (RNA-seq) confirmed that pancreatic hormone transcripts were significantly higher (>2 fold difference; p<0.05) in the adult pancreas, whereas *Hes1*, a negative regulator of the pro-endocrine gene *Neurog3* (also known as *Ngn3^23^*) was expressed at significantly higher levels in gallbladder cells (**Fig. 1E**). In line with the embryonic data (**Fig. 1B**), the expression of *Pdx1* was similar (**Fig 1E**) across adult gallbladders and the pancreas. Validation of RNA-seq data using real-time qPCR, confirmed similar *Pdx1* expression in both adult tissues (**Fig. 1F**). Relatively lower levels of *Neurog3* transcripts in adult gallbladder presented with a significantly higher level of *Hes1*. Pancreatic islets and gallbladder epithelial cells of *Pdx1^GFP/w^* mice had insulin- as well as Pdx1-coexpressing cells (**Fig. 1G**). Using another (MIP-GFP) mouse model^24^, we confirmed mouse *Ins1* promoter activity in adult mouse gallbladder cells (**Fig. S1D**). Immune-electron microscopy of mouse gallbladder epithelial cells confirmed the presence (albeit lower number) of electron-dense insulin secretory vesicles (**Fig. 1H**) within gallbladder epithelium. These studies demonstrate pancreatic endocrine gene expression in embryonic and adult mouse gallbladder epithelium. Moreover, we also demonstrate the capacity of adult mouse gallbladder epithelium to translate and package insulin for secretion.

**Fig. 1:**
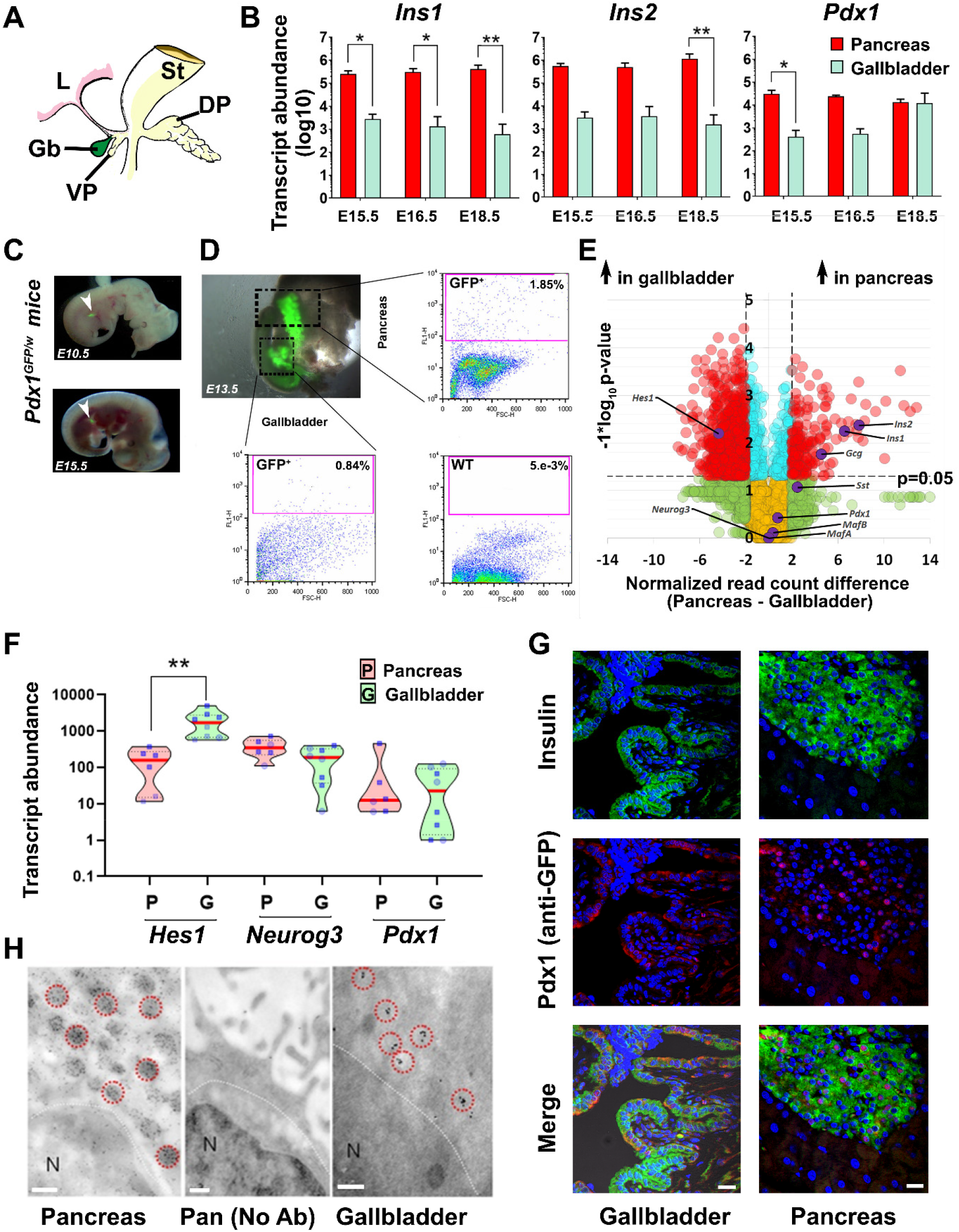
Mouse gallbladder and pancreas development. (**A**) Mouse pancreatic gut schematic displaying developing stomach (St), Liver (L), dorsal pancreatic bud (DP), and the ventral pancreatic bud (VP) that gives rise to the gallbladder (Gb) and ventral pancreas. (**B**) TaqMan®-based real-time quantitative PCR for rodent insulin genes (*Ins1, Ins2*) and the master regulatory transcription factor (*Pdx1*) in developing mouse pancreas and gallbladder tissues harvested at embryonic day (*E*) 15.5, 16.5 and 18.5. Data represent mean+SD from three different litters of FvB/NJ mice, each with at least 6-7 embryos/litter. Transcript abundance was analyzed using two-way ANOVA with Sidak’s adjustment for multiple comparisons. (**C, D**) *Pdx1^GFP/w^* reporter mouse embryos were obtained from timed pregnancies (E10.5, E13.5, and E15.5) and GFP fluorescence was used to identify pancreatic buds. Representative flow cytometry plots of pancreas and gallbladder tissues from *Pdx1^GFP/w^* (GFP^+^) and *Pdx1^w/w^* (wild type WT) mice are presented with proportions of GFP^+^ cells indicated. Acquisition and gating parameters were identical across WT and GFP^+^ tissues. Experiments were repeated at least three times with tissues pooled from 6-7 embryos. (**E**) Volcano plot for bulk RNA-sequencing data from the adult mouse gallbladder (*n*=6) and pancreas (*n*=6). Normalized read count difference between pancreas and gallbladder is plotted on X-axis and statistical significance (−log10 p-value), is presented on Y-axis. The dashed horizontal line represents p-value=0.05, whereas dashed vertical lines represent a 2-fold normalized DEseq value difference. Significantly altered (p<0.05 and >2-fold different) transcripts are presented in red. Selected important pancreatic genes on the volcano plot are labelled and highlighted in purple. (**F**) Real-time qPCR data from postnatal mouse pancreas and gallbladder (n=6-10 animals) for *Hes1*, *Neurog3,* and *Pdx1* gene transcripts analyzed using Kruskal-Wallis with Dunn’s multiple comparisons test. Each dot within the violins represents a different sample (circles: age group 1-7 days; squares: age group 3-8 weeks). The horizontal solid red line within each polygon represents the median, the horizontal black dotted line represents quartiles, and the polygons represent the density of individual data points and extend to min/max values. (**G**) Immunostaining of insulin (green) and Pdx1/GFP (red) in the pancreatic islets and gallbladder epithelial cells of *Pdx1^GFP/w^* reporter mice. Nuclei (DNA) are shown in blue. The scale bar is 20 μm. (**H**) Electron microscopy images from the same CD1 mouse pancreatic islet and gallbladder epithelial cells following immuno-gold labelling of insulin granules. Data were validated in four different preparations. Immuno-gold labelled insulin vesicles are indicated with red circles. N denotes the nucleus (all panels) while the middle panel represents a no-antibody pancreatic control. The scale bar is 0.2 μm. *=p<0.05; **=p<0.01 and ****=p<0.0001.

### Pancreatic endocrine gene expression in developing human gallbladder

We extended our mouse studies to developing human pancreas and gallbladders accessible from 11 fetuses at early (16-20 weeks gestational age; WGA; n=5) or late stages (>20WGA; n=6, **Fig 2A**) of pregnancy. In comparison with pancreatic tissue, all the islet hormones (*INS, GCG, SST*) were expressed at significantly lower abundance (p<0.001; **Fig 2B**) in gallbladder cells. Differences between islet and gallbladder (pro-)hormone transcript levels were significant across early (16-20WGA; p<0.01, n=5, **Fig 2B**) but not later stages (**Fig 2B**). Histological (**Fig. 2C,D**) and confocal microscopy analysis (**Fig. 2E,F**) of >20WGA human pancreas and gallbladders confirmed tissue-specific morphology and presence of hormone-containing cells. We then compared the expression of pancreas-enriched transcription factors and receptors between four pairs of the gallbladder(G) and pancreas(P) samples from human fetal donors (**Fig. 2G**). We also validated the expression of selected pancreatic genes in remaining human pancreatic and gallbladder fetal tissues. Unlike mouse development, the significantly lower (p=0.04) level of human *NEUROG3* in developing gallbladder cells was not associated with any differences in *HES1* expression (**Fig. 2H**). Although levels of *GCK* gene transcripts were significantly higher in the developing pancreas (p=0.02; **Fig. 2I**), *GLP1R* and the glucose transporter *GLUT2* (also known as *SLCA2*) were similar across these developing tissues (**Fig. 2I**). Several other genes (*CDH1, HB9, NEUROD1, PAX6,* and *PCSK2*) were expressed at significantly higher levels in the developing human pancreas as compared to their levels in gallbladders (**Fig. S2**). No significant differences were observed across the expression of zinc finger transcription factors (*GATA4, GATA6*, **Fig. 2H**), islet integrin αV (*ITGAV*) and the histone deacetylases (*HDAC1-3*, **Fig. S2**).

**Fig. 2:**
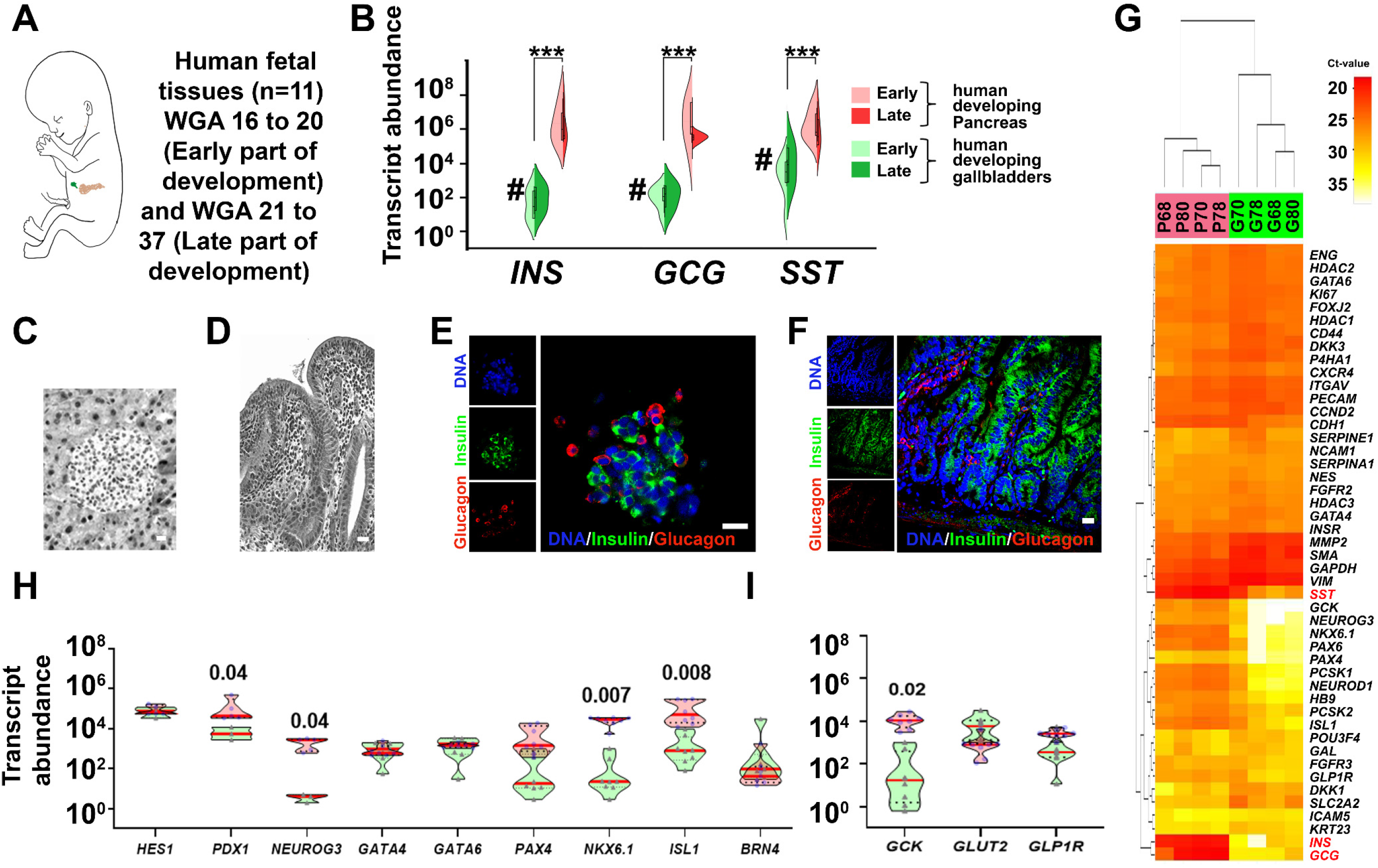
Human gallbladders exhibit several markers of pancreatic endocrine lineage during embryonic development. (**A**) Schematic of developing human fetus. Pancreas and gallbladder tissues were available from a total of 11 different fetuses, which were classified as early (>20 weeks gestation age/WGA; n=5) or late (>20 WGA; n=6) development. (**B**) TaqMan^®^-based real-time qPCR for key pancreatic endocrine hormones (*INS, GCG,* and *SST*) in developing human pancreas and gallbladder tissues are plotted based on their developmental stage. Split violin plots compared gene expression in early vs late gallbladders (shades of green) and pancreas (shades of red). The horizontal line within each bar of the split-violins represents the median, bars extend to quartiles, and the polygons represent the density of data points and extend to min/max values. Data were analyzed using Kruskal-Wallis with Dunn’s multiple comparisons tests; ***p<0.001 denotes significant difference between all pancreas and all gallbladder samples, whilst **#** p<0.01, denotes a significant difference between the early pancreas and early gallbladder samples. (**C, D**) Representative hematoxylin and eosin (H&E)-stained images of human fetal pancreatic islet and gallbladder sections (in greyscale). (**E, F**) Immunostaining of insulin (green) and glucagon (red) in the human pancreatic islet and gallbladder epithelial cells from late development. Nuclei (DNA) are shown in blue. The scale bar for C-F is 20 μm. (**G**) An unsupervised bidirectional hierarchical plot of 48 important pancreatic genes and transcription factors in the same human fetal pancreas and gallbladder tissues (n=4; indicated by the sample number) using Euclidean distance metric and average linkage is presented. Heat map represents normalized qPCR Ct-values (coloured bar) of each gene (gene symbol listed on the right vertical axis) with low Ct-values/high expression in orange-red colour and higher Ct-values/low expression in shades of yellow to white. (**H, I**) Real-time qPCR data from the human fetal pancreas (n=5-11) and gallbladder (n=3-11) for pancreatic genes and transcription factors, analyzed using two-way ANOVA with Fisher’s LSD test. The horizontal solid red line within each polygon represents the median, the horizontal black dotted line represents quartiles, and the polygons represent the density of data points extending to the min/max lues.

### Similarities in the developing human pancreas and gallbladders are retained in adult life

Bright-field and confocal microscopy of adult human pancreatic islets and gallbladder epithelial cells confirmed that the morphology and islet-specific protein production observed in fetal stages is retained in adult human gallbladder cells (**Fig. 3A**). Adult human gallbladder epithelial cells isolated as epithelial sheets (see SOM) form hollow spheres of epithelial cells, which are immuno-positive for C-peptide and PDX1 (**Fig. 3A**), confirming the capacity of the gallbladder epithelial cells to process the prohormone into mature insulin and C-peptide. Similar to human pancreatic islets^17^, gallbladder-derived epithelial cell clusters show immuno-positivity for E-cadherin and β-catenin (**Fig. 3A**). Bulk RNA sequencing in freshly isolated adult human islets and gallbladder epithelial cells confirmed a large number of genes that are significantly different across these functionally diverse tissues (**Fig. 3B**). RNA-seq coverage maps confirm that adult human gallbladder transcripts for *INS, GCG, SST* and *PDX1* mapped to the same genomic regions (**Fig. 3C**) as in islets. Validation in a different set of the freshly isolated adult human islet and gallbladder samples using TaqMan-based real-time qPCR (**Fig 3D-E, S3A**), confirmed that islet hormones were at significantly higher (several 100-thousand-fold) levels in human islets, relative to those in gallbladder epithelium, while the expression of *GLUT2* and insulin receptor (*INSR*) were comparable (**Fig 3D**). Pancreatic transcription factors (including *PDX1 and MAFA*) but not *NEUROG3*, were significantly more abundant in pancreatic islets (**Fig. 3E**). We compared our human gallbladder RNA-seq data with a publicly available RNA expression dataset of human laser capture microdissected β-cells from non-diabetic individuals (E-GEOD-20966)^25^. Interestingly, several gene transcripts had similar expression levels across these two functionally diverse tissues (**Fig. 3F, Table S5**). Gene ontology (GO) analysis of gallbladder transcripts filtered for β-cell expression indicated several GO-terms that are relevant to insulin packaging and secretion (**Fig 3G**), cellular development/differentiation (**Fig S3B**), vesicle transport (**Fig S3C**), mitochondrial structure, and function (**Fig S3D**) as well as carbohydrate metabolism including glucose metabolic processes (**Fig S3E**). We then mapped the expression of 21 highly abundant human gallbladder-enriched gene transcripts identified through our RNA-seq datasets (**Fig S4A**) to publicly available pancreatic single-cell(sc)RNA-sequencing datasets (**Fig. S4B**). Intriguingly, 17 of the 21 gene transcripts were present in acinar and/or ductal cells, while ~50% (10 of 21) of the gallbladder-enriched gene transcripts were also transcribed by pancreatic α- and/or β-cells (**Fig S4C**). In summary, our data demonstrate that the inherent capacity of human gallbladder epithelial cells for pro-endocrine gene transcript and protein expression, is retained in the post-natal stage and that the adult human α- and β-cells transcribe several of the gallbladder-enriched gene transcripts.

**Fig. 3:**
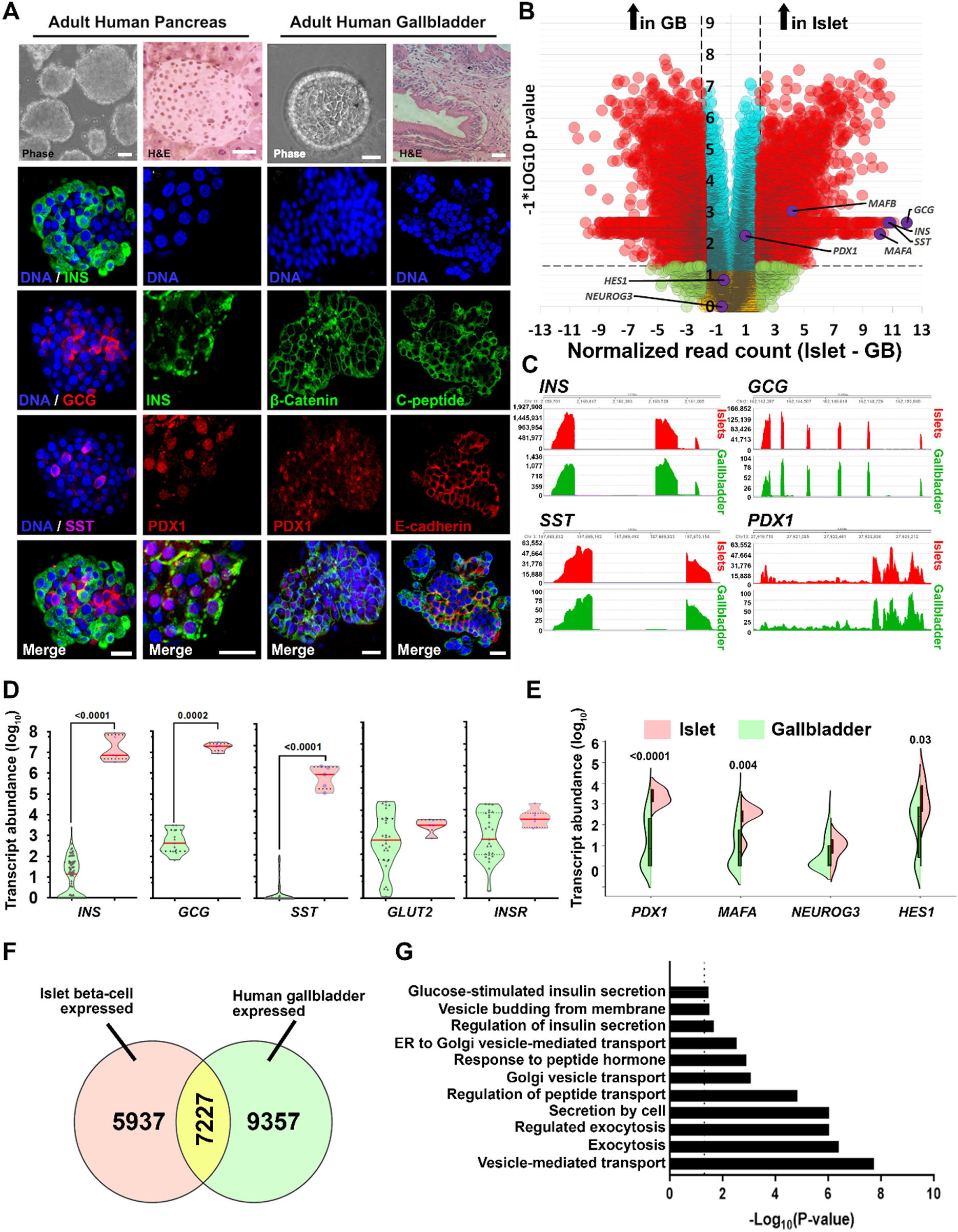
Adult human gallbladder cells express low levels of pancreatic islet hormones. (**A**) Representative phase-contrast, as well as hematoxylin and eosin (H&E) images of adult human pancreatic islets and gallbladder epithelium, are shown in the top panel. Confocal images for insulin/C-peptide, glucagon, somatostatin, PDX1, β-catenin, or E-cadherin immunostaining are presented. Nuclei (DNA) are shown in blue. The scale bar is 20 μm. (**B**) Volcano plot for bulk RNA-sequencing data from the adult human gallbladder (n=7) and islet samples (n =6). Normalized read count difference between islet and gallbladder are plotted on the X-axis and statistical significance (−log10 p-value) is on Y-axis. The dashed horizontal line represents a p-value=0.05, whereas the dashed vertical lines represent a 2-fold difference. Significantly altered (p<0.05 and >2-fold difference) transcripts are presented in red. Important pancreatic genes are labelled and highlighted in purple. (**C**) Coverage plots (obtained from bulk RNA sequencing; panel **B**) for *INS, GCG, SST,* and *PDX1* expression across their gene regions (shown at the top of each plot) in human islets and gallbladder. Y-axis presents read counts. (**D, E**) Real-time PCR data from human islets (n=5-6) and gallbladder (n=27-98) for pancreatic genes, analyzed using the Mann-Whitney test. Each dot in the violin plot (**D**) represents a different sample. The horizontal solid red line within each polygon represents the median, the horizontal black dotted line represents quartiles, and the polygons represent the density of data points, extending to min/max values. The horizontal line within each bar of the split-violins (**E**) represents the median, bars represent quartiles, and the polygons present the density of data points, extending to min/max values. The exact p-value for significant difference between pancreas and gallbladder are presented. (**F**) Venn diagram demonstrating the number of genes common between our bulk gallbladder RNA sequencing (n=6) and publicly available dataset of human islet-beta cells (see methods). (**G**) The most significant and relevant gene ontology terms enriched in gallbladder cells and associated with endocrine pancreatic β-cell function are presented. The X-axis represents −log10 p-value, the dashed vertical line represents the significant p-value=0.05 and relevant pathways are provided on the Y-axis.

### Adult human gallbladder epithelial cells can proliferate and differentiate *in vitro*

We established a unique protocol to isolate gallbladder epithelium without enzymatic digestion, via gentle scrapping of gallbladder epithelial cells using a sterile scalpel blade (**Fig 4A).** The epithelial sheets obtained by scraping, close on to themselves to form hollow, epithelial clusters of cells (**Fig 4B**), which retain the capacity to migrate and proliferate *in vitro* as mesenchymal-like cells (**Fig 4C**) through a process believed to involve epithelial-to-mesenchymal transition (EMT; **Figure 4D**). Using an unbiased, dual-thymidine analogue-based cell lineage tracing technique^1 26^, we detected the presence of CldU^+^, IdU^+,^ and C-peptide^+^ cells (**Fig. 4E**) in gallbladder-derived cells, propagating under optimal cell culture conditions (**Fig S5A**). Eventually, all proliferating populations of gallbladder epithelial cells acquire a mesenchymal-like phenotype and begin expressing mesenchymal gene transcripts (**Fig. S5B**) and proteins (**Fig 4F**), including typical surface antigens expressed by mesenchymal stem cells (**Fig S5C**). We then assessed the chromatin landscape at the insulin gene in freshly isolated human islets and gallbladder clusters. At two different sites of insulin gene (−275 and +1318), human islets displayed open chromatin conformation as confirmed by higher levels of H3H4Ac and H3K9Ac; histone modifications associated with active genes along with lower levels of H3K9-trimethylation; associated with silenced/inactive genes. In the freshly isolated gallbladder epithelial cells, these chromatin modifications were not significantly different from those observed in adult human islets (**Fig 4G**). Analysis of pancreatic gene promoters in passage 5 gallbladder-derived mesenchymal cells demonstrated a more inactive chromatin conformation (H3K9-Me3 and H3K9-Me2) at insulin and Neurogenin 3 promoters, whereas an ‘open’/accessible chromatin conformation (H3K4-Me2) at the *HES1* gene promoter (**Fig 4H**). Interestingly, the *PDX1* gene promoter retained an active/open promoter conformation in these cells (**Fig 4H**). These data corroborate with the lack of insulin expression in gallbladder-derived mesenchymal cells and support the potential for re-expression of insulin gene transcripts through chromatin conformational changes or via repressing *HES1* expression. We, therefore, assessed endocrine differentiation of gallbladder-derived mesenchymal cells using small molecules that are known to be DNA methyltransferase inhibitors or histone deacetylase inhibitors^27–29^ and/or via forced expression of a dominant-negative (Δ)*HES1*, or of the pro-endocrine transcription factors (*PDX1, MAFA, NEUROG3*). Although differentiation to insulin-expressing islet-like clusters was observed over 14 days (**Fig 4I**), none of the small molecules assessed induced significantly higher insulin expression compared to vehicle controls (**Fig 4J**). On the other hand, we obtained a consistent and significant increase in *INS* transcript expression using both (Δ*HES1* and pro-endocrine transcription factor) overexpression strategies (**Fig. 4K**). Our data support the view that gallbladder-/biliary duct-cells can be differentiated to promote insulin expression.

**Fig. 4:**
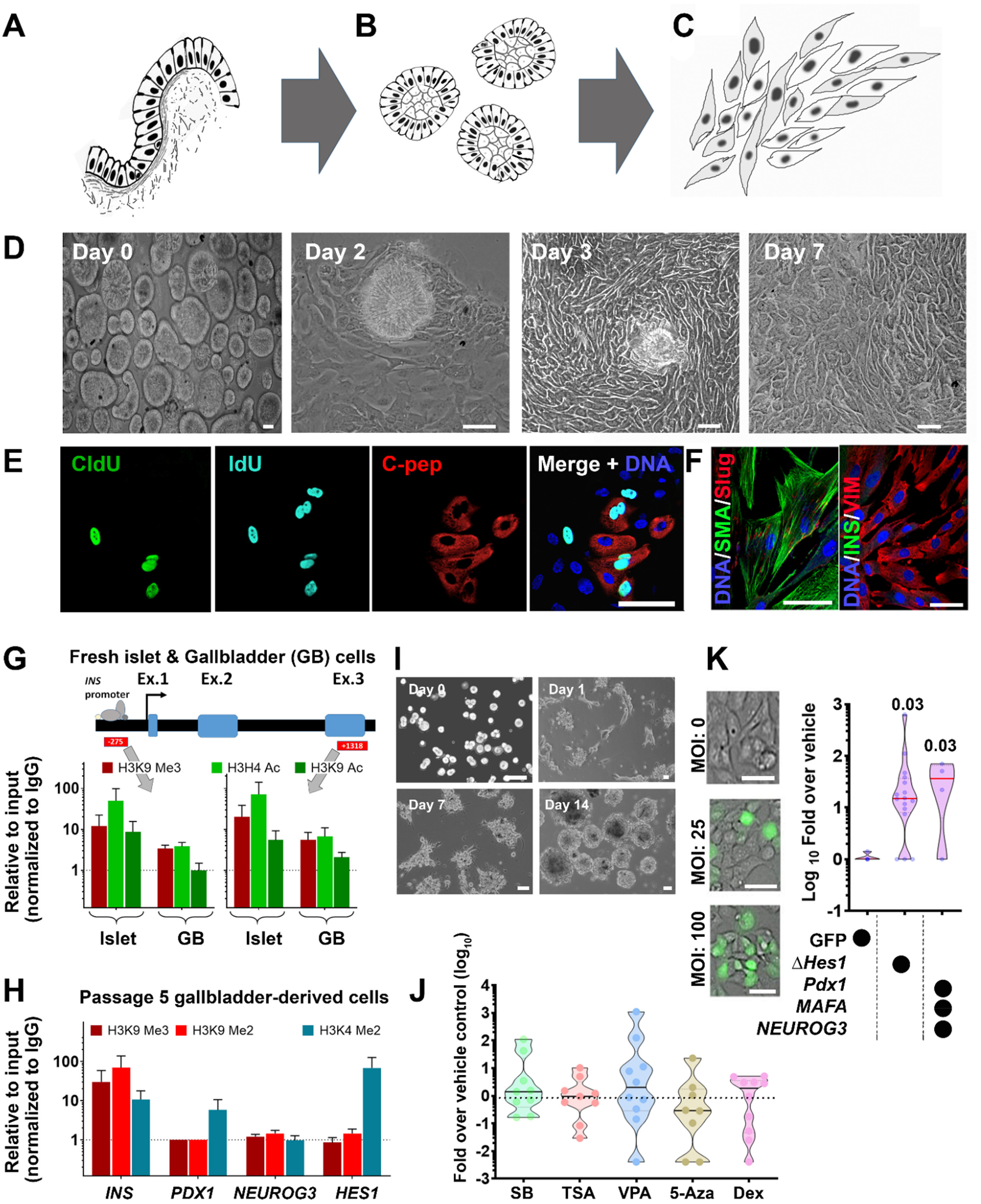
Adult human gallbladder-derived cells can be induced to differentiate *in vitro*. (**A-C**) A schematic of isolating and propagating adult human gallbladder epithelial cells *in vitro*. The inner epithelial cell lining of the gallbladder (**A**) is scraped off with a sterile scalpel. The resulting epithelial sheet curls up into clusters (**B**) that then adhere to culture plates and proliferate/propagate as mesenchymal cells (**C**). (**D**) Representative phase-contrast images of the process described through panels A-C; epithelial clusters from adult human gallbladders on day 0, attach (day 2), migrate out (day 3) and continue to grow/proliferate (day 7) *in vitro*. (**E**) Immunostaining of CldU (green), IdU (aqua), and C-peptide (red) in cultures of gallbladder epithelial cells after 7 days involving 3-day exposure to CldU, 1 day of wash-out and next 3 days of incubation in IdU-containing medium. Nuclei (DNA) are shown in blue as a representative from n=10 preparations. (**F**) Proliferating human gallbladder-derived cells express mesenchymal proteins *in vitro*. Nuclei (DNA) are shown in blue. (**G**) One site (−275) in the insulin promoter region and another (+1318) in the exon 3 region were assessed for the presence of H3K9 Me3 (inactive mark) as well as H3H4 Ac and H3K9 Ac (active transcription marks) using chromatin immunoprecipitation assay in freshly isolated human islets (n=6) and gallbladder epithelial cells (n=3-5). (**H**) Promoter regions of insulin, *PDX1*, *NEUROG3* and *HES1* were assessed for the presence of H3K9 Me3 and H3K9 Me2 (inactive marks) as well as H3K4 Me2 (active transcription mark) using chromatin immunoprecipitation assay in passage 5 gallbladder-derived mesenchymal cells (mean+SEM; n=2-6 different biological preparations). Data in **G**and **H** are presented relative to input and normalized to IgG. The dotted line represents expression in isotype control (IgG) samples. (**I**) Representative phase-contrast images of the differentiation process involving trypsinization (day 0), reaggregation (day 1), and cluster formation (day 7-day14). (**J**) Real-time qPCR data for insulin gene in differentiated gallbladder cells using five different DNMT and HDAC inhibitors/hypomethylation agents (SB: sodium butyrate; TSA: Trichostatin A; VPA: Valproic acid; 5-Aza: 5-Aza-2′-deoxycytidine; Dex: Dexamethasone). Data are presented as fold-over vehicle control from 9-10 different biological samples/experiments. (**K**) Phase-contrast images of transduction efficiency using GFP expressing adenoviral vectors. Real-time qPCR data for insulin gene in differentiated gallbladder cells using a vehicle (GFP alone) or a dominant-negative *HES1* or a combination of *PDX1*, *NEUROG3,* and *MAFA*. Data are presented as fold-over vehicle control from 4-15 different biological samples and analyzed using the Kruskal-Wallis test. The exact p-value after adjusting for multiple comparisons are presented. Each dot in the violin plots (in **J** and **K**) represents a different sample. The horizontal solid line within each polygon represents the median, the horizontal black dotted line represents quartiles, and the polygons represent the density of individual data points and extend to min/max values. The scale bar is 20 μm in all panels.

### Functional analyses of insulin-producing human gallbladder cells

We compared genes known to be important in insulin sensing and exocytosis using bulk RNA-seq data of adult human islet and gallbladder epithelial cells. Although gene transcripts of islet hormones and the glucose sensor (*GCK*) were significantly lower in gallbladder epithelial cells, the transcripts of the two glucose transporters were either similar (for *SLC2A1*) or significantly higher (for *SLC2A2*) in adult gallbladder epithelial cells (**Fig. 5A**) than those in adult human islets. Except for pancreatic ATP-sensitive K+ channel *ABCC8* (or *SUR1*) and the transmembrane protein Synaptotagmin-7 (*SYT7*), gallbladder epithelial cells and human islets contained similar levels (**Fig. 5B**) of gene transcripts for *KCNJ11* (or *KIR6.2*), the L-type Calcium channel *CACNA1C* (encoding CAV1.3 protein), the islet Syntaxins and binding proteins (*STX1A, STX3,* and *STXBP2*), the synaptosomal-associated protein (*SNAP25*) and gene transcripts encoding the vesicle-associated membrane proteins (*VAMP2* and *VAMP8*). These comparisons (**Fig. 5B, 3G)** indicate that gallbladder epithelial cells express the set of genes necessary for insulin exocytosis. Freshly isolated gallbladder epithelial cells (**Fig. 5C**) contain insulin protein and also release insulin/C-peptide in response to glucose (**Fig. 5D**) indicating successful processing and secretion of insulin following glucose exposure *in vitro*. We then transplanted freshly isolated adult human gallbladder epithelial cells under the kidney capsule of immuno-compromised (NOD/SCID) mice (**Fig. 5E**). A functional assessment carried on day 30 using intraperitoneal glucose injection resulted in detectable levels of human insulin in mouse circulation (**Fig. 5F**). To understand *in situ* insulin secretion from human gallbladder cells, we collected blood from the median cubital vein (peripheral) and cystic vein (gallbladder) at 30 minutes following intravenous glucose injection from individuals, who were enrolled for cholecystectomy. Intriguingly, insulin levels in the cystic vein were higher than those in the cubital vein for 80% (n=4 of 5) individuals assessed in this study (**Fig. 5G**). Overall, we demonstrate that naturally occurring insulin-secreting cells within the gallbladder have relevant insulin secretory machinery and can respond to physiological changes in glucose concentrations.

**Fig. 5:**
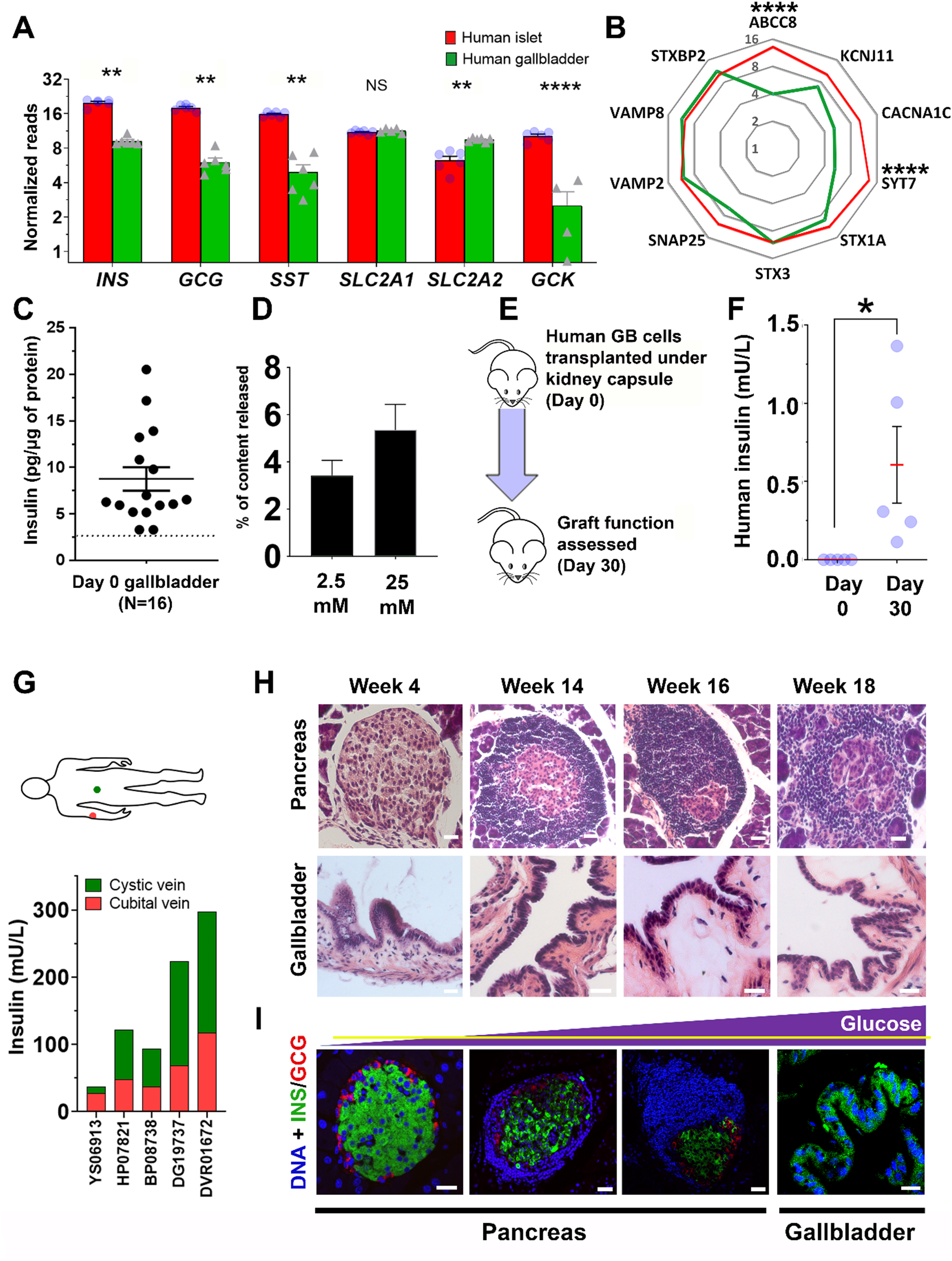
Gallbladder epithelium contains functional glucose-responsive cells. (**A**) Expression of key pancreatic hormones, glucose transporters, and glucose-sensing enzyme gene transcripts in adult human islets (n=6) and adult human gallbladder (n=4-6) samples. Data are obtained from bulk RNA sequencing and presented as normalized read counts. Bar graphs present mean+SEM and differences between groups are assessed using two-tailed t-tests. (**B**) Expression of secretory function-related genes in human islets (n=6) and gallbladder (n=7) is presented as normalized mean read counts in radar plot (red line: pancreas, and green line: gallbladder). Data analyzed using the Kruskal-Wallis test followed by adjustment for multiple comparisons using Dunn’s test. (**C**) Insulin content in freshly isolated human gallbladder epithelial cells. Data are obtained from n=16 different biological samples. Each dot in the scatter plot indicates a separate sample with lines indicating mean+SEM. (**D**) C-peptide release in response to two different concentrations of glucose is presented as % of total content. Data are from 7-8 different freshly isolated human gallbladder cells and presented as mean+SEM. (**E**) Human gallbladder epithelial cells were isolated and transplanted in NOD/SCID mice and functional assessment was carried out on day 0 and day 30. (**F**) Human insulin was measured in mouse serum. Data are obtained from n=5 different gallbladder samples transplanted in mice. Each dot in the scatter plot represents a different sample and the red line and error bars represent mean+SEM. Significance was calculated using Welch’s t-test. (**G**) Circulating insulin concentrations in cystic (green) or median cubital (red) veins were measured in 5 individuals (labeled on X-axis). Stacked bars represent insulin concentration from cystic (gallbladder; green) and peripheral (cubital; red) veins. (**H**) Representative hematoxylin and eosin (H&E) images of pancreas and gallbladder sections obtained from NOD mice of different ages. (**I**) Immunostaining of insulin (green) and glucagon (red) in islet and gallbladder sections obtained from NOD mice at different ages with increasing concentration of circulating glucose (yellow horizontal line depicting normoglycemic levels in the purple triangle representing increasing glucose concentrations). Nuclei (DNA) are shown in blue. The scale bar is 20 μm in all panels. *p<0.05; *** p<0.001 and ****p<0.0001.

### Insulin-producing gallbladder epithelial cells in diabetes

To understand if insulin-containing gallbladder epithelial cells also elicit immune-mediated attack during diabetes progression, we examined immune infiltration in gallbladder and islets from female NOD mice. Since NOD females show significant infiltration at 12-14 weeks of age, we obtained gallbladder and pancreas from NOD mice at 4-, 14-, 16- and 18-weeks of age (**Fig. 5H**). Predictably, the pancreas from these mice demonstrated increasing infiltration of immune cells (**Fig. 5H**) at 14-18 weeks of age, but no infiltration was observed in their gallbladders. Immunostaining for islet hormones confirmed the decreasing number (quantitative data not shown) of insulin-producing cells in the pancreas as plasma glucose increased (**Fig. 5I**). Insulin-immunoreactive cells were present in gallbladder epithelial cells even when NOD mice showed high glucose levels at 18 weeks (**Fig. 5I**), indicating that gallbladder epithelial cells can potentially escape immune recognition during type 1 diabetes progression in NOD mice.

## Discussion

The origin/proximity of the gallbladder endoderm to the developing pancreas is evolutionarily conserved across lower vertebrates^30 31^, and mammalian biliary duct cells are shown to produce islet hormones^1 7 32–35^. We confirm that human and mouse gallbladder epithelial cells contain functional insulin-containing cells during development and retain them in adult stages. Our studies in NOD mice, which naturally develop immune-mediated T1D, indicate that gallbladder epithelial cells do not demonstrate immune infiltration in T1D and that these cells may continue to produce and release insulin. This interesting observation could be due to different autoimmune antigenic repertoire between islet β-cells and gallbladder insulin-producing cells. Analysis of splice variants in gallbladder (GSE152419; n=7) and human islet (GSE152111; n=66 and GSE134068; n=18) RNA-sequencing datasets indicate high differential splice index for human insulin variant (ENST00000250971) in islets (splicing index 0.14, n=66 islets or 0.17, n=18 islets, respectively) compared to no differential splicing observed in human gallbladder epithelial cells (ENST00000250971, splicing index 0.0, n=7). Genetic or environmental factors leading to Endoplasmic Reticulum (ER) stress are of critical importance in regulating the expression of alternative splice variants via alternative reading frames of insulin^36 37^. Metabolic or inflammatory stress may lead to the generation of novel splice forms via recognition of newly created splice site^38^ and the generation of defective ribosomal peptides (DRiPs)^39 40^. Such DRiPs may act as neoantigens to which central immune tolerance is absent. The lower abundance of insulin, as well as the lack of alternate insulin splice forms in the gallbladder, may explain the lack of autoimmune cells in the gallbladder of diabetic NOD mice. Another possibility is high levels of the taurine-conjugated bile acid Tauroursodeoxycholic Acid (TUDCA) in the gallbladder, which is known to reduce ER stress and protect islet β-cells in T1D mouse models^41 42^. TUDCA is currently in clinical trials for T1D (NCT02218619) and further studies understanding the potential role of ER stress and DRiPs in the gallbladder during T1D progression are merited. Interestingly, all cases of neonatal diabetes with undetectable levels of immunoreactive-insulin also report gallbladder aplasia/hypoplasia^43^, signifying the common developmental plans for these two functionally diverse organs. Although the pancreas of individuals with long-standing type 1 diabetes may continue to produce insulin^44^, understanding the contribution of insulin from gallbladder epithelial cells would help in advancing preventive or therapeutic interventions in T1D.

The present study has several strengths. This is the first study collectively demonstrating the inter-species and age-related similarities across two functionally diverse, but developmentally related organs; the gallbladder and the pancreas. For the first time, a comparison between the gallbladder and pancreatic cells/samples using multiple techniques to demonstrate similarities in chromatin conformation (ChIP-PCR), gene transcription (bulk/scRNA sequencing, TaqMan qPCR, gene reporter analyses), insulin production (confocal microscopy, ELISA, immune-electron microscopy) and insulin release (animal models, human cells and clinical *in situ* measurements) is presented. Direct analysis of insulin-producing cells in gallbladder and pancreas from NOD mice lays important emphasis on the therapeutic and preventative mechanisms that may be important in consideration for future research in T1D.

A limitation is that our studies do not explain the possible mechanisms underlying the biological variation observed in insulin transcript abundance across different samples. Future studies would need to assess proinsulin and insulin ratios in the same gallbladder epithelial cell samples to understand the concentrations of processed and mature insulin and be statistically powered to understand if differences are related to ethnic, genetic, or environmental components. Secondly, although the differentiation of gallbladder-derived cells was not the principal aim of this study, differentiation strategies need improvement to enhance insulin production in gallbladder-derived progenitor cells.

In summary, we demonstrate the inherent property of mouse and human gallbladder cells to transcribe, translate, package, and release insulin in response to glucose. The capacity for insulin gene transcription in gallbladder epithelium appears to be driven by the developmental similarity, which programs pancreatic islet epigenetic and transcriptomic signatures in the gallbladder. The demonstration that this intrinsic insulin production escapes autoimmune damage during T1D progression renews interest in this functionally diverse organ.

## General

Authors acknowledge and thank the support from surgical teams, consenting donors or families of the donors as well as infrastructure support obtained through the Rebecca L Cooper Foundation (to AAH) and provided by the Faculty of Medicine & Health, University of Sydney. AAH, HET, AHE and EGS acknowledge the assistance from Dr. Andrew M. Holland and Dr. Suzanne J. Micallef in the provision of the *Pdx1^GFP/w^* reporter mice and Prof. Manami Hara for the MIP-GFP mice. AAH and DT acknowledge the support from Prof. Jonathan Slack and Prof. Harry Heimberg for the viral vectors used in the study. Support from Dr. Ramesh Dumbre, Dr. Pabitra Sahoo, Ms. Fahmida Khan, Ms. Sophie Breedveld, RPA pathology services, Bosch facilities (BMSF and LCAF) at the University of Sydney is also acknowledged.

## Author contributions

Lab work, data acquisition and analysis: MVJ, SS, WKMW, SNS, CXD, RJF, MW, PP, GJ, SNY, YC, NH, SM, GVR and AAH. Resource development and provision/clinical studies: Dhan T, GVR, DNR, HET, TL, WJH, AGE, VMJ, EGS, DM, Data analysis and interpretation: MVJ, SS, SNS, WKMW, David T, LTD, AAH. Research study planning: AAH. Writing: MVJ, WKMW, SS, LTD and AAH. All authors read and acknowledge the final draft for submission.

## Data and materials availability

The Biospecimen Reporting for Improved Study Quality (BRISQ) guidelines^45^ were followed. Single-cell sequencing (Panc8) datasets (n=14,890) were extracted from already available datasets (GSE84133, GSE85241, E-MTAB-5061, GSE83139, GSE81608). Our data on bulk RNA sequencing for the human gallbladder dataset (GSE152419; n=7) and human islet datasets (n=84 islet samples through GSE152111, n=66 and GSE134068, n=18), are uploaded to Gene Expression Omnibus (GEO) database and will be made publicly available with this publication. Data for mouse gallbladder and pancreas bulk RNA-sequencing are also available through GSE152419.

