## Supplementary methods, tables and figures for "A pro-endocrine pancreatic transcriptional program established during development is retained in human gallbladder epithelial cells"

***Running Title: Insulin-producing cells in human gallbladder***


**Mugdha V. Joglekar**[1,°], **Subhshri Sahu**[1,+,°], **Wilson KM Wong**[1,°], Sarang N. Satoor[1], Charlotte X. Dong[1], Ryan J Farr[1], Michael D. Williams[1], Prapti Pandya[1], Gaurang Jhala[2], Sundy N.Y. Yang[1], Yi Vee Chew[3], Nicola Hetherington[1], Dhan Thiruchevlam[4], Sasikala Mitnala[5], Guduru V Rao[5], Duvvuru Nageshwar Reddy[5], Thomas Loudovaris[2], Wayne J. Hawthorne[3], Andrew G. Elefanty[6], Vinay M. Joglekar[7], Edouard G. Stanley[6], David Martin[8], Helen E. Thomas[2], David Tosh[9], Louise T. Dalgaard[10,] and **Anandwardhan A. Hardikar**[1,*]

[1] Diabetes and Islet Biology Group, School of Medicine, Western Sydney University, 30 Narellan Road & Gilchrist Drive, Campbelltown, NSW 2560, Australia.
[2] Immunology and Diabetes Group, St. Vincent's Institute for Medical Research, 9 Princes St, Fitzroy, VIC 3065, Australia
[3] Westmead Clinical School and the Westmead Millenium Institute, 176 Hawkesbury Rd, University of Sydney, Westmead, NSW 2145, Australia
[4] Department of Gastroenterology, St. Vincent's Hospital, Melbourne, VIC 3065 Australia
[5] Surgical Gastroenterology Research, Asian Institute of Gastroenterology, Hyderabad, India.
[6] Stem Cell Technology division, Murdoch Childrens Research Institute, Flemington Road, Parkville, VIC 3052, Australia
[7] Shree Seva Medical Foundation, Off Pune -Bangalore NH4, Shirwal, MH412801, India
[8] Upper GI Surgery, Strathfield Hospital, 2/3 Everton Rd, Strathfield, NSW 2135, Australia
Current address: National Center for Cell Science, Ganeshkhind Road, Pune 411007, India.
[9] Department of Biology and Biochemistry, University of Bath, 4 South 0.66, Claverton Down, Bath BA2 7AY, United Kingdom
[10] Section for Eukaryotic Cell Biology, Department of Science and Environment, Roskilde University, Roskilde, Denmark

° These authors contributed equally
[+] Previously at Diabetes and Islet Biology Group, National Center for Cell Science, Ganeshkhind, Pune, 411007, India.

[*] Address all correspondence to:
Anandwardhan A. Hardikar, PhD
Diabetes and Islet Biology Group, School of Medicine, Western Sydney University,
30.2.27 Goldsmith & David Pilgrim Ave, Campbelltown, NSW 2560, AUSTRALIA

Web: https://www.isletbiology.info/

**Supplementary Online Materials**

**Detailed methods**

**Materials and Methods**

**Animals:** FVB/NJ mice were used for embryonic gene expression and insulin content, CD1 mice were used for immune-gold labelling studies at the University of Bath, C57BL/6J mice were used for RNA sequencing, qPCR or ChIP studies, Pdx1-GFP, MIP-GFP and NOD mice were used for gene reporter and immunostaining studies. All animals were maintained at the experimental animal facilities in India, UK, or Australia according to guidelines outlined by the respective institute's animal care and use committee. Ethical approvals for the study were obtained from animal ethics committees at the National Centre for Cell Science in India, St. Vincent's Hospital in Melbourne, and the University of Sydney. Breeding pairs were set, and pregnancy was confirmed by observing vaginal smears. Pregnant females and newborn mice were euthanized at pre-defined intervals and pancreatic buds or pancreas, as well as gallbladder tissue, was carefully dissected without cross-contamination using a stereomicroscope. Tissue samples at each of these time points were used for RNA isolation (in Trizol) and immunostaining (in 4% freshly prepared paraformaldehyde).

**Human tissue collection:** Adult gallbladder, pancreas, and islets samples were obtained following human research ethics committee (HREC) approvals from Sydney Local Health District and St. Vincent's Hospital in Melbourne. Fetal gallbladder and pancreas were obtained following informed consent from prospective parents who consented to fetal tissue for research following elective termination of pregnancy (<20 WGA) or following late abortions/miscarriage (>20WGA) as per human research ethics committee (HREC) approvals from Shree Seva Medical Foundation, India and National Centre for Cell Science, India. Human cadaveric non-diabetic pancreas and islet samples were obtained as part of the research consented tissues through the Australian Islet Transplantation Program (at Westmead Hospital,

Sydney, and the St Vincent's Institute, Melbourne). Human gallbladders were obtained as surgical waste tissues (non-cancerous) after cholecystectomies from the surgical teams (at Strathfield Private Hospital, Royal Prince Alfred Hospital, Sydney and The St. Vincent's Hospital, Melbourne and National Centre for Cell Science, Pune, India). Tissue samples were stored for RNA isolation (in Trizol), immunostaining (in 4% freshly prepared paraformaldehyde) or processed for cell culture as detailed below. For in situ glucose stimulation study, we consented five individuals undergoing cholecystectomies for blood collection at the Asian Institute of Gastroenterology. Blood was collected before and after the glucose challenge. The study was performed according to the Declaration of Helsinki II. Study participants gave informed written consent before inclusion as per the ethical committee approval from the Asian Institute of Gastroenterology, India.

**Gallbladder cell culture:** Gallbladder tissues were collected in transport medium (M199 with 25mM HEPES and 2mM glutamine medium with 2X antibiotics) and processed in the lab. The gallbladder sample was washed thoroughly with the serum-free medium (M199 with 25mM HEPES and 2mM glutamine + Hams F12k) containing 100U/mL penicillin and 100μg/mL streptomycin (Gibco, Carlsbad, CA) and fungizone (Gibco, Carlsbad, CA) until bile was removed. The inner surface of the gallbladder was then gently scraped off with a disposable, aseptic scalpel blade to isolate the epithelial lining. These isolated cells were washed with the wash medium (described above), centrifuged at 1000g for 2 minutes and the cell pellet was re-suspended and plated in serum-containing medium (10% FBS + M199 with 25mM HEPES and 2mM glutamine + Hams F12k) with penicillin, streptomycin (Gibco, Carlsbad, CA). This medium is hereafter referred to as a growth-promoting medium or serum-containing medium (SCM). Cells were maintained in an incubator at 37°C and with humidified 5% CO2 in the air and passaged 1:2 when confluent using trypsin (Gibco, Carlsbad, CA) + EDTA. MTT assay was used to understand the media composition supporting the maximum viability and metabolic activity of isolated cells. Briefly, 50,000 cells were seeded in each well of 96-well plates and allowed to adhere overnight. Different media and serum concentrations were formulated and added to cells (24 conditions) in quadruplets. After 24h and 48h, MTT reagent (Sigma-Aldrich, St Louis, MO) was added to each well and after 2-3 hours, conversion of MTT to formazan was measured using a spectrophotometer. The experiment was repeated with 3-4 different gallbladder preparations.

**In vitro differentiation of human adult gallbladder-derived cells:** Human adult gallbladder-derived mesenchymal-like cells are trypsinized and differentiated following an optimized protocol in our lab [1-3]. Briefly, these cells are plated on day (D) 0 in serum-free medium (D0 SFM) that consists of DMEM: F12 + 1% Bovine serum albumin (BSA) +1X Insulin-Transferrin-Selenium (ITS). Single cells on day 0 start to aggregate into islet-like clusters (ICAs). On day 1, the medium was changed with D0 SFM to remove dead cells. On day 4 and day 7, ICAs were exposed to a D4 medium that contains 0.3mM taurine + D0 SFM. On day 10, ICAs are exposed to D10 SFM that contains nicotinamide (1mM) and exendin-4 (100nM) along with D4 SFM. On D14 these ICAs were harvested for RNA isolation. In some experiments, DNMT and HDAC inhibitors were added during the entire time of differentiation at the following final concentrations: 1 mM Sodium butyrate (SB), 100nM Trichostatin A (TSA), 1mM Valproic acid (VPA), 2µM 5-aza-2'-deoxycytidine (5-Aza) and 1µM dexamethasone (dex). Vehicle control (PBS or DMSO) was used for comparisons. Adenoviral vectors for *PDX1*, *NEUROG3,* and *MAFA* were used at two different multiplicity of infection (MOI) in gallbladder-derived cells. Transductions were performed as described earlier[4]. Transduced cells were differentiated following the protocol detailed above and GFP-alone adenovirus transduced cells were used for gene expression comparisons.

**RNA isolation, cDNA synthesis, and quantitative real-time PCR:** RNA isolation from fresh human and mouse tissues at different stages (embryonic, adult) was carried out using Trizol

(Invitrogen, Carlsbad, CA). RNA quality and quantity were measured on ND-2000 spectrophotometer (NanoDrop Technologies, Wilmington, DE). cDNA synthesis was carried out using High Capacity cDNA Reverse Transcription Kit (Thermo Fisher Scientific, Foster City, CA) as per manufacturer's recommendations. Quantitative real-time PCR was performed with TaqMan primer and probes mix (Thermo Fisher Scientific, Foster City, CA) for genes listed in **Table S2** and TaqMan Fast Universal PCR Master Mix (Thermo Fisher Scientific, Foster City, CA) in 5µL reaction in 96 well optical clear plates. The cycle threshold (Ct) values of all the genes were normalized to the housekeeping gene 18s rRNA. Transcript abundance was calculated using normalized Ct-values and the formula "transcript abundance=$2^{(39\text{-Ct value})}$", where '39' is the limit of detection on the real-time PCR system[5]. TaqMan Low-Density Array (TLDA) cards were designed for selected pancreatic genes and housekeeping genes (**Table S3**) and obtained from Thermo Fisher Scientific. TLDA cards were used to assess gene expression in human fetal pancreas and gallbladder tissues using the manufacturer's protocol standardized for TLDA cards on the 7900 HT system (Thermo Fisher Scientific, Foster City, CA). Normalized Ct-values were used to plot hierarchical cluster heatmaps. A customized OpenArray™ Human mRNA panel (Thermo Fisher Scientific, Waltham, MA) was designed for analyzing the 45 selected pancreatic gene transcripts and housekeeping control in adult human islets and gallbladder (**Table S4**). The customized panels were used following the manufacturer's protocol standardized for gene expression using OpenArray on QuantStudioTM 12K Flex Real-Time PCR platform (Thermo Fisher Scientific, Waltham, MA). Normalized Ct-values were used to plot hierarchical cluster heatmaps.

**Bulk RNA sequencing:** RNA sequencing on mouse tissues was performed as detailed in [6] using Ion Total RNA-Seq Kit v2, Ion OneTouch 200 Template Kit v2 DL (Thermo Fisher Scientific, Waltham, MA) and sequenced on Ion Torrent PGM Instrument using Ion PGM 316™ Chip and Ion PGM 200 Sequencing Kit v2 (Thermo Fisher Scientific, Waltham, MA).

Adult human pancreatic islets and gallbladder epithelial cell samples (gallbladder dataset GSE152419; n=7 and human islet dataset GSE152111, n=66) were sequenced as 150 paired-end reads on the HiSeq4000 platform as detailed elsewhere[7]. Poly-T oligo-attached magnetic beads were used to purify the mRNA from total RNA for these samples.

**Bulk RNA-Seq analysis:** The Strand Next-generation sequencing (NGS) version 2.5 software was used to analyze the RNA-Seq data. For mouse samples, reads were aligned to the mouse mmu10 (UCSC) transcriptome and genome together with novel splice variants, using the Ensembl genes and transcript model. While for human samples, reads were aligned to the human hg38 transcriptome and genome together (with novel splice variants) using the Ensembl genes and transcript model. Raw reads were aligned with a minimum of 90% identity, a maximum of 5% gaps, and a minimum match length of 25 base pairs. The output of read-pairs with more than five valid matches were not reported. Quality trimming was applied to trim 3' end with an average quality less than 10 and on a poorly aligned portion at 3' end post alignment. Following alignment, raw reads were filtered to remove reads with an average base quality below 20 and reads which have failed quality control (QC). For mouse samples, an average of 2.57 million clean reads per sample were aligned to the reference transcriptome/genome (with novel splice variants; GSE152419). For human samples, an average of 29 million clean reads were generated through bulk RNA-seq of gallbladder epithelial cells (GSE152419; n=7) and human islets (GSE152111, n=66) and 78 million clean reads for another set of human islet samples (GSE134068, n=18). DEseq was used to quantify and normalize the aligned reads, with the threshold normalized count set at 1[8]. Baseline transformation was not applied to the pre-processing of the aligned read input data.

**Pancreatic single-cell (sc)RNA sequencing analysis:** Panc8 single-cell sequencing data (n=14,890) was extracted from public datasets (GSE84133, GSE85241, E-MTAB-5061, GSE83139, GSE81608). It was analyzed via R studio version 1.2.5033 (built under the R 3.6.1) using the SeuratData (version 0.2.1), Seurat, ggplot2 and cowplot packages. The original Seurat Panc8 package contains eight different pancreas single-cell RNA-sequencing datasets from across five technologies (inDrop, CEL-Seq1, CEL-Seq2, Smart-Seq2, and Fluidigm C1). To improve the integrity of the data, low read count datasets obtained through the inDrop technology were excluded. Analytical workflow included data preprocessing and feature selection, dimension reduction, and identification of "anchor" correspondences between datasets, filtering, scoring, and weighting of anchor correspondences, and data matrix correction, or data transfer across experiments as described elsewhere [9 10].

**Immunostaining:** For immunostaining, tissue sections or freshly isolated epithelial clusters, islets or cultured gallbladder cells were fixed in 4% PFA. Tissues were embedded in paraffin and 5-10µm sections were placed on slides for further processing. Hematoxylin and Eosin staining were done on tissue sections using standard protocol after deparaffinization. For immunochemistry, permeabilization was done in chilled methanol (50% v/v in water) or 0.5% triton X 100 followed by blocking with 4% normal donkey serum (NDS, Sigma-Aldrich, St Louis, MO). Incubation with the primary antibody was at a dilution of 1:100 or 1:200 overnight at 4°C. Cells were then washed 5 times with 1X PBS containing $Ca^{2+}$ and $Mg^{2+}$ (Gibco, Carlsbad, CA) and incubated with secondary antibody at 37°C. Cells were then thoroughly washed and mounted in Vectashield mountant (Vector Laboratories, Peterborough, UK) containing Hoechst 33342 (Invitrogen, Carlsbad, CA). Primary antibodies used were rabbit polyclonal antibody to human C-peptide and guinea pig anti-Insulin (both from Linco Research Inc, St. Charles, MO), rabbit anti-Somatostatin (Dako, Santa Barbara, CA), mouse anti-Glucagon, mouse anti-GFP, mouse anti-smooth muscle actin and rabbit anti-slug (all from Sigma-Aldrich, St. Louis, MO), goat anti-Pdx1 (Abcam, Cambridge, UK), mouse anti-E-cadherin and mouse anti beta-catenin (BD Biosciences, Franklin Lakes, NJ), and mouse monoclonal anti-Vimentin (Chemicon Int. Inc, Temecula, CA). Alexa-Fluor 488, 546, and 633 secondary antibodies (Invitrogen, Carlsbad, CA) were used at 1:200 dilution. Images were scanned and assessed using a Zeiss LSM 510 laser confocal microscope after setting their thresholds below saturation. Laser power and other parameters were identical for all samples.

**Immuno-electron microscopy:** CD1 mouse gallbladder and pancreas isolated from the same animal were fixed in 0.5% Glutaraldehyde, 4% Paraformaldehyde (Agar Scientifics, England) and 2.5mM $CaCl_2$ (Thermo Fisher) in 0.1M Sodium cacodylate (Agar Scientifics, England) buffer (SCB) at 4°C overnight, rinsed next day in a solution of 0.1M SCB and 2.5mM $CaCl_2$ for 10 minutes with four changes at room temperature. Samples were dehydrated in ethanol (70%. 90%, 100% dry and 100% dry) for 15 minutes at room temperature, infiltrated with LR white (Agar Scientifics, England) for 1h at room temperature, overnight at 4°C and again at room temperature for 1h. Each tissue was transferred to a gelatin capsule (size 0; Agar Scientifics, England) and the capsule was filled completely with LR white resin and allowed to polymerize at 50°C for 24 hours. 100nm sections were cut and mounted on formvar –carbon-coated nickel slot grid (Agar Scientifics, England) and immunolabelled by floating the mounted grids on respective solutions. Sample (grids) were exposed to 0.05M glycine (Sigma, St. Louis, MO) in PBS to block aldehyde sites for 15 minutes, rinsed briefly with PBS, blocked in blocking solution with 10% normal goat serum, 0.5% BSA in PBS for 15 minutes, rinsed briefly and then incubated in guinea pig anti-Insulin (Linco Research Inc, St. Charles, MO) at a dilution of 1:500 overnight at 4°C. Grids were then washed with PBS, 3 times for 10 minutes each, and incubated in secondary antibodies (gold tagged goat anti guinea pig IgG, British Biolcell, Cardiff, UK) at 1:50 dilution for 2 hours at room temperature covered, washed in 0.1M SCB for 3 times. The grids were then post-fixed with 1% glutaraldehyde in 0.1M SCB for 15 minutes, rinsed twice with distilled water for 5 minutes each, and then dried. The grids were then stained in 2% Uranyl Acetate for 10 minutes, covered with lead citrate for approximately 4 minutes in NaOH containing chamber, then washed with fresh distilled water, dried and then imaged using transmission electron microscopy (Jeol 1200EX TEM, University of Bath, UK).

**Flow cytometry:** Pdx1-GFP and MIP-GFP mice were euthanized at predetermined embryonic and adult stages. Pancreas and gallbladder tissues were stored on ice, finely chopped followed by collagenase digestion (3mg/mL, 10 min at 37°C) to generate a single-cell suspension. Cells were strained through a 70µm cell strainer and then resuspended in FACS buffer (Ca2$^{+}$, $Mg^{2+}$ free PBS +2% FCS) and acquired on BD FACSCalibur™ (BD Biosciences, Franklin Lakes, NJ). Gallbladder-derived mesenchymal-like monolayer cells at passage 5 were harvested using trypsin. Cells were washed twice with FACS buffer, blocked with 5% BSA in 1X PBS (Ca2$^{+}$ and $Mg^{2+}$ free) for 30 minutes, incubated with fluorescently-conjugated antibodies for 1h, washed twice with FACS buffer, fixed with 4% PFA and then acquired using FACSCalibur™. The antibodies used were Phycoerythrin (PE) labelled Rat IgG2 (isotype control), anti-CD29, anti-CD44, anti-CD90, and anti-CD105 (all PE-labelled BD Biosciences, Franklin Lakes, NJ) at dilutions recommended by the manufacturer. The data were analyzed using CellQuest Pro software. Isotype control or wild type tissues were used to set up gating. PI$^{+}$ (dead) cells were excluded from analyses. All acquisition parameters were kept unchanged for the same type of tissue or cells.

**Cell lineage tracing:** To follow the propagation of the insulin-positive cells *in vitro*, we used two thymidine analogues CldU and IdU (Sigma-Aldrich, St Louis, MO) as described before[11]. Freshly isolated gallbladder epithelial cells were seeded on Lab-Tek chamber slides (Nunc, Rochester, NY 14625 USA). CldU was added to freshly isolated cells at a concentration of 10□M and was incubated in a growth-promoting medium for 3 days, followed by one day of wash (without any analogue) and again a 3-day pulse of IdU at a concentration of 10□M. On day 7 the cells in culture were fixed in 4% PFA, washed once with PBS and then 0.2% Triton X 100 was added for 5min at room temperature. Antigen retrieval was carried out by microwaving the slides in 0.01M sodium citrate, pH 6.0 for 6 min. at high power until the citrate buffer started boiling and then kept for 20 minutes in the same citrate buffer at low power in the microwave. After cooling down to room temperature, cells were washed once with 1X PBS containing $Ca^{2+}$ and $Mg^{2+}$ and then kept in 1.5N HCL (diluted in 1X PBS containing $Ca^{2+}$ and $Mg^{2+}$) for the next 40 minutes at room temperature. This is then followed by blocking with 4% NDS at RT for 30 min. The rest of the steps involve immunostaining procedure as described above. Immunostaining was carried out using antibodies for rat anti-CldU (Accurate Chemicals, Westbury, NY), mouse anti-IdU (BD biosciences, Franklin Lakes, NJ) and rabbit anti-C-peptide (Dako, Santa Barbara, CA) at a concentration of 1:100. Alexa-Fluor 488, 546, and 633 secondary antibodies (Invitrogen, Carlsbad, CA) were used at 1:200 dilutions.

**Chromatin immunoprecipitation (ChIP):** Adult mouse gallbladder and pancreas, freshly isolated epithelial cells from the adult human gallbladder, freshly isolated human islets, and gallbladder-derived mesenchymal cells in culture at passage 5 were used in ChIP assay as described before[12]. Briefly, cells were cross-linked, washed with buffers containing protease inhibitor cocktail (PIC, Sigma-Aldrich, St Louis, MO) and sonicated in lysis buffer to generate DNA fragments of 200-400 base-pairs. Chromatins were immunoprecipitated using 2µg of specific dimethyl and trimethyl antibodies for H3K4 and H3K9 as well as for acetylation of H3K9 and H3 and H4 (Millipore, Billerica, MA). Precipitation cocktails included protein A/G plus beads (Pierce, Pittsburgh, PA), sonicated salmon sperm DNA (Amersham Biosciences Pittsburgh, PA), and BSA (USB corporations, Cleveland, OH USA). Rabbit and mouse IgG (Upstate, Millipore, Billerica, MA) were used as isotype controls. Chromatin was eluted using 2% SDS, 0.1M NaHCO3 and 10mM DTT. Cross-links were reversed by incubating the eluted chromatin in 4M NaCl overnight at 65°C. This was followed by proteinase-K digestion and DNA extraction using phenol-chloroform- isoamyl alcohol.  Input, immunoprecipitated and isotype control DNA was resuspended in nuclease-free water and used for SYBR green or TaqMan qPCR with primers listed in **Table S5** and with Fast SYBR green or TaqMan™ Fast master mix (Thermo Fisher Scientific, Waltham, MA) on the ViiA7™ Real-Time PCR System platform (Thermo Fisher Scientific, Waltham, MA).

**Glucose stimulated insulin secretion:** Gallbladder epithelial clusters were handpicked under a phase-contrast microscope. These were then washed with Krebs-Ringer Bicarbonate HEPES (KRBH) buffer containing 0.1% BSA (Sigma, St Louis, MO) and exposed in quadruplicates to basal (2.5mM Glucose) or stimulated (25mM Glucose) buffer, for 1 hour at 37°C. At the end of the incubation/exposure cells were settled down or centrifuged at 300g for 1 min to pellet. The supernatant was collected and assayed for insulin /C-peptide. The insulin content in the tissues/cells was measured by sonicating them in 200-500µL of acid ethanol, depending upon the size of a tissue or cell pellet. Total protein concentration was measured using Bradford assay (for mouse samples) or ThermoFisher Scientific's Qubit protein assay (for human samples). Insulin or C-peptide concentrations were measured by ELISA kit (Mercodia, Winston Salem, NC). Circulating human insulin was measured using the same kits (Mercodia) following plasma separation. Plasma was separated by centrifugation at 1000g for 10 min and stored at -80°C before performing insulin ELISAs. Blood glucose was measured using the Accu-Chek glucometer.

**Transplantation of human gallbladder epithelial cells:** Transplantation of freshly isolated gallbladder epithelial cells was carried out on 8-12 wk old male NOD/SCID to assess their function in response to glucose stimulation as described before[2]. During this procedure, animals were placed on their back after anaesthesia with isoflurane. A total of 500-700 freshly isolated gallbladder epithelial clusters were added into ~20µL of animal blood (obtained from the tail) to form a blood clot, which was then transplanted under the kidney capsule of NOD/SCID mice without losing any of the cell clusters. Animals were opened by making the left lateral incision to expose the left kidney. The kidney was gently pulled out and a superficial cut was made in the kidney capsule. The blood clot containing gallbladder cells was placed below the kidney capsule. The capsule was then gently massaged to close the cut. The kidney was then placed back to its original position. The incision was closed using 3-4 absorbable sutures (Davis-Geck, Manati, PR) and an autoclip wound clipper (BD biosciences, Franklin Lakes, NJ). Topical ointment (Soframycin®) was applied over the sutured wounds following surgery and animals were administered analgesics (Buprenorphine 0.05mg/kg every 12 hours for 3 days) as and when necessary. On day 30 of the surgery, blood was collected at 30 minutes following a 2g/kg body weight glucose load.

**Pathway analysis:** To analyze enrichment for β-cell pathways lists of gallbladder expressed genes (GSE152419) were compared with β-cell-expressed genes (from E-GEOD-20966) using GO-analysis on Pantherdb.org[13]. Pre-analytic workflows included cleaning up entries not mapping to protein-coding gene symbols. Gene lists of 16584 gallbladder transcripts and 13164 β-cell transcripts were compared for overlaps in transcripts using Venn diagrams (https://bioinfogp.cnb.csic.es/tools/venny/index.html), identifying 7227 genes to be present in both gallbladder and β-cell expression data set. GO-analysis was performed using the list of 7227 gallbladder transcripts also present in β-cells and compared to the list of all human genes as reference. Statistical overrepresentation was calculated using Fisher's exact test, using Bonferroni correction for multiple testing.

**Statistical analysis:** Statistical analyses were performed using GraphPad Prism 8.4.1 (GraphPad Software, San Diego, CA, USA) or the R-software (ver. 3.6.2, R Foundation for Statistical Computing, Vienna, Austria), SPSS Statistics 27 (Chicago, IL, USA) or Microsoft Excel (ver. 2016; Microsoft, Redmond, WA, USA). R software was used to perform unsupervised hierarchical clustering maps using heatmap.2 function in gplots. Add-on and other R packages XLconnect (with ActivePerl software) and RColorBrewer were used for dataset import and visualization along with R package gplots. GraphPad Prism was used to perform all remaining analyses using appropriate statistical tests and corrected for multiple comparisons if required. Details of each statistical test, number of replicates and the number of animals/biological preparations are provided in the respective figure legends. Split volcano and spider/radar plots were created using BioVinci data visualization package. Kolmogorov-Smirnov test was used to check for data normality in SPSS. F-test was performed to check for variance in Excel. For non-normally distributed data, the two-tailed Mann-Whitney test was used to calculate the P-value with no ties computed as performed in R. While for normally distributed data with equal or unequal variance, two-tailed Student's or Welch's t-test was used to calculate the P-value in Excel, respectively.

**Supplementary Figures**

**Fig. S1: Similarity and differences in mouse embryonic pancreatic and gallbladder cells**

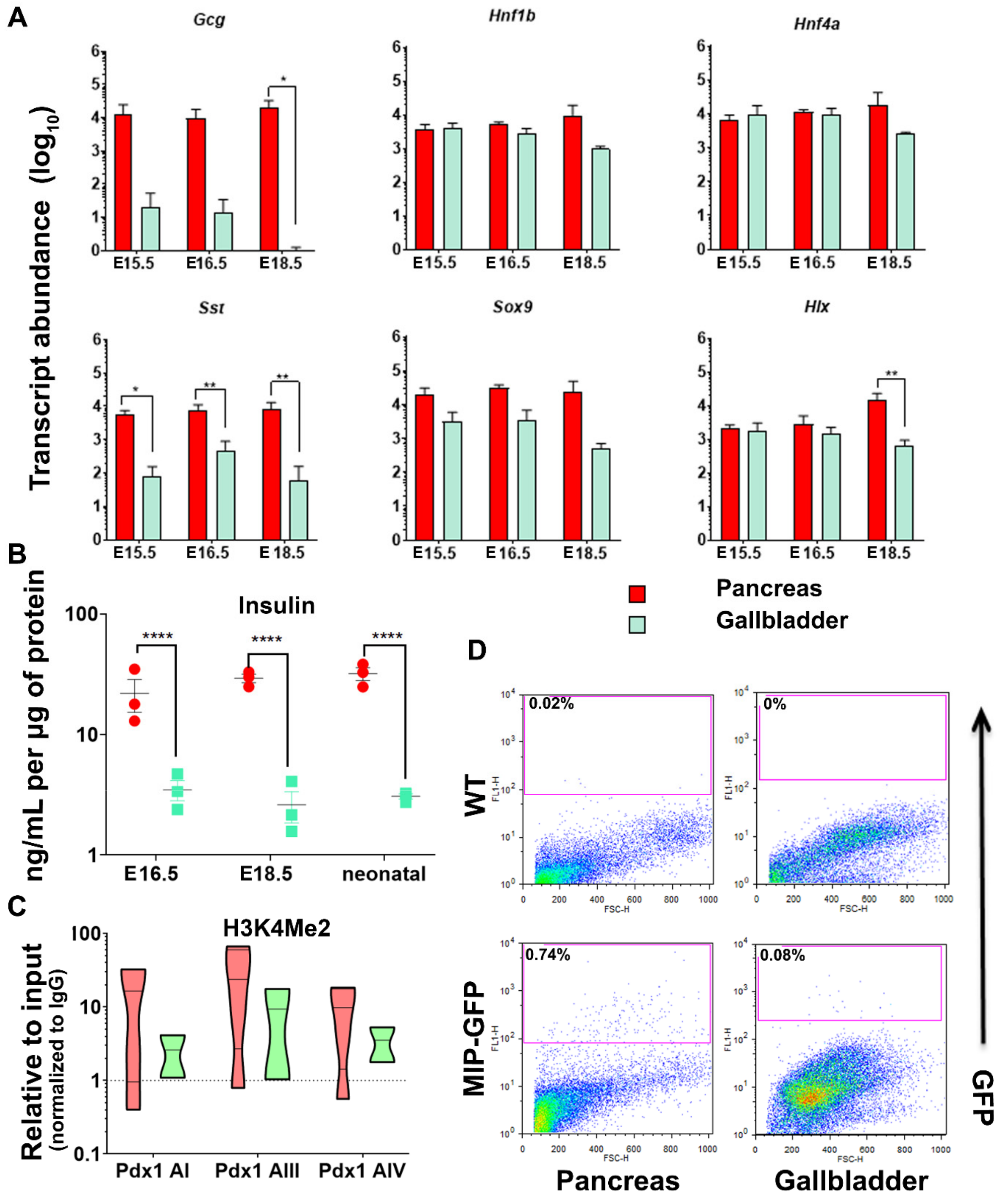

***Fig. S1: Related to Fig. 1:***

(**A**) TaqMan®-based real-time qPCR for mouse pancreatic genes in developing pancreas and gallbladder tissues harvested at embryonic day (E) 15.5, E16.5, and E18.5. Data are obtained from three different litters of FvB/NJ mice, each with at least 6-7 embryos/litter. Transcript abundance was calculated as described in methods and data were analyzed using two-way ANOVA with Sidak's multiple comparisons test. (**B**) Insulin content in developing mouse pancreas and gallbladder tissues harvested at E16.5, E18.5, and neonatal day 1 pups. Data are obtained from pooled tissues from three different litters of FvB/NJ mice, each with at least 6-7 embryos/pups and presented after normalizing to total protein. Aligned dot plots present mean$\pm$SEM. Significance is calculated using two-way ANOVA with Fisher's LSD test. (**C**) Three sites in the promoter region of mouse *Pdx1* were assessed for the presence of H3K4 Me2 (active transcription mark) using chromatin immunoprecipitation assay in adult C57BL/6J mice pancreas (n=4) and gallbladders (n=2 experiments with >6 mice gallbladders /experiment). Data are plotted relative to input and normalized to IgG. The dotted line represents the expression levels for isotype control (IgG) samples. The horizontal solid red line within each violin plot represents the median, and the polygons represent the density of data points and extend to min/max values. (**D**) Representative flow cytometry plots of pancreas and gallbladder tissues from adult (6-10 week old) *MIP-GFP* reporter and wild type (WT) mice. The percentage of $GFP^+$ cells is indicated for each plot. Acquisition and gating parameters were identical across wild type and $GFP^+$ cells. Experiments were repeated at least three times with tissues pooled from 3-5 animals at each time. Throughout the figure, bars/spots/violins for the pancreas are in red and for the gallbladder in green. *=$p<0.05$; **=$p<0.01$ and ****=$p<0.0001$.

1 **Fig. S2: Similarity and differences in developing human pancreatic and gallbladder cells**

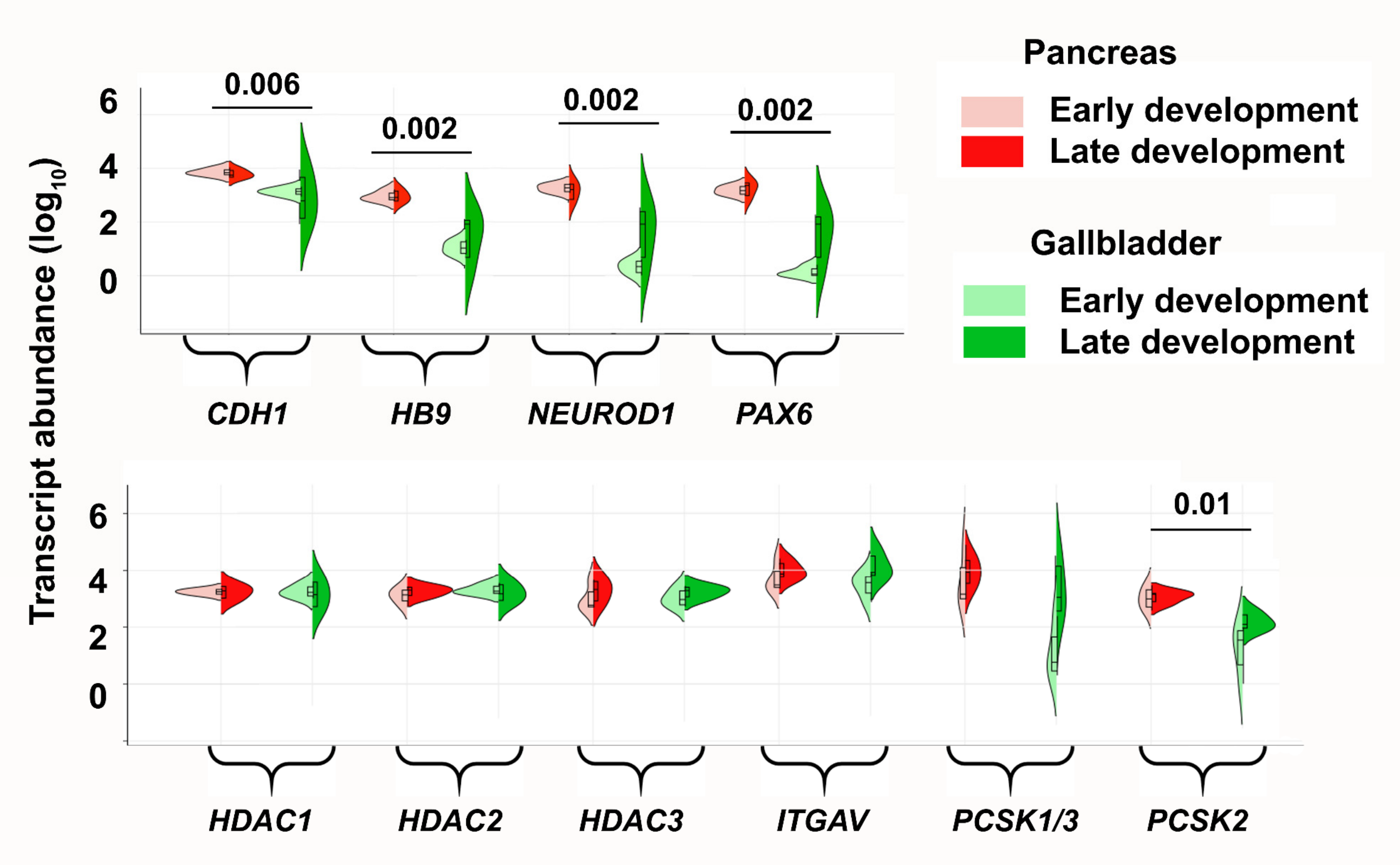


2

***<u>Fig. S2: Related to Fig. 2:</u>***

Pancreas and gallbladder tissues from different (n=11) human fetuses were classified to early (<20 weeks gestation age/WGA; n=5) or late (>20 WGA; n=6) development. TaqMan®-based real-time qPCR for pancreatic/epithelial gene transcripts and histone deacetylation enzymes in developing human pancreas and gallbladder tissues are presented and analyzed using Kruskal-Wallis with Dunn's multiple comparisons test. Split violin plots compared the gene expression in early vs late gallbladders (shades of green) and pancreas (shades of red). The horizontal line within each bar of the split-violins represents the median, bars extend to quartiles, and the polygons represent the density of data points and extend to min/max values. The p-values represent a two-tailed Welch's test between all (early and late) gallbladder vs. pancreas samples. No statistically significant differences were observed within the developmental groups (early vs. late), for either of the pancreatic or gallbladder tissue samples for any of the gene transcripts shown.

**Fig. S3: Gene expression pathways enriched in adult human gallbladder epithelial cells**

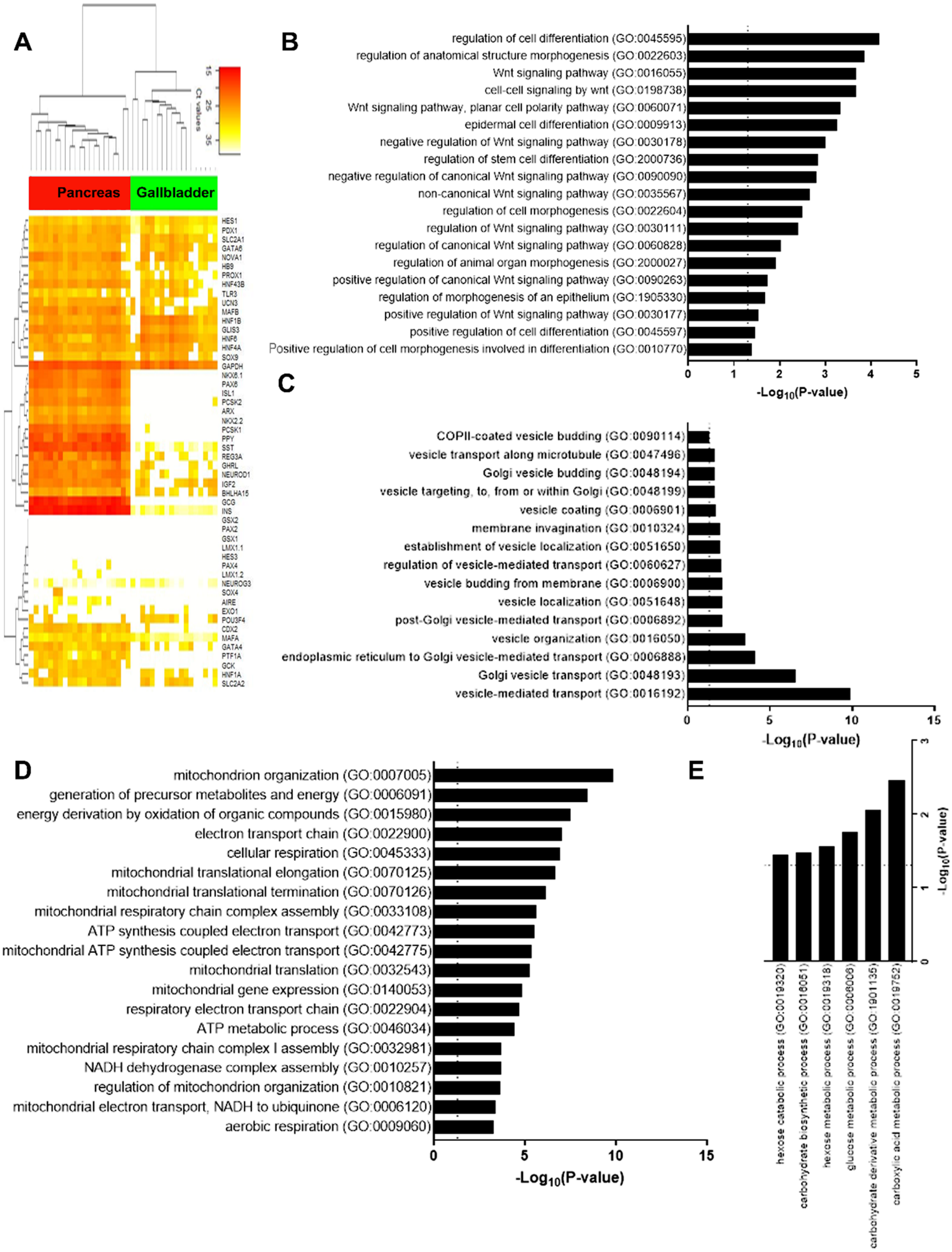

***<u>Fig. S3: Related to Fig. 3:</u>***

(**A**) An unsupervised bidirectional hierarchical cluster of 51 genes known to be associated with or necessary for normal pancreas development/function, profiled in adult human pancreatic islets (red, n=21) and human gallbladder epithelial cells (green, n=18) was plotted using Euclidean distance metric and average linkage. The heat map representing normalized qPCR Ct-values (colour bar) for each gene (listed on right Y-axis) with low Ct-values/high expression in orange-red colour and higher Ct-values/low expression in shades of yellow to white. Data consist of measurement on ViiA7 (*INS*, *GCG*, *MAFA*, *NEUROG3*, *HES1,* and *PDX1*) or using a custom TaqMan OpenArray platform. (**B-E**) The most significant and relevant Biological Process Gene Ontology (GO) categories significantly enriched in transcripts from gallbladder cells filtered for beta-cell expression. Data obtained from RNA sequencing was used to identify the gallbladder expressed genes and total gallbladder transcripts were filtered for beta cell-expressed transcripts obtained from E-GEOD-20966. The X-axis represents −log10 p-value, the dotted vertical line represents the significant p-value=0.05 and relevant pathways are provided on the Y-axis. Pathways involved in development and differentiation (**B**), vesicle transport (**C**), mitochondrial function (**D**), and carbohydrate metabolism (**E**) are presented.

**Fig.S4: Human pancreatic scRNA-Seq reveals expression of gallbladder-enriched genes**

A

| Gene ID | Gene Symbol | Average read count |
|---|---|---|
| ENSG00000120885 | CLU | 88253.3 |
| ENSG00000162896 | PIGR | 87702.2 |
| ENSG00000184009 | ACTG1 | 33362.5 |
| ENSG00000245532 | NEAT1 | 31933.0 |
| ENSG00000251562 | MALAT1 | 30391.9 |
| ENSG00000145321 | GC | 29773.6 |
| ENSG00000169894 | MUC3A | 29571.2 |
| ENSG00000198848 | CES1 | 25759.7 |
| ENSG00000171345 | KRT19 | 21751.4 |
| ENSG00000149131 | SERPING1 | 20698.7 |
| ENSG00000119547 | ONECUT2 | 19692.2 |
| ENSG00000117289 | TXNIP | 17374.4 |
| ENSG00000066230 | SLC9A3 | 17234.0 |
| ENSG00000064651 | SLC12A2 | 15914.2 |
| ENSG00000185499 | MUC1 | 15872.7 |
| ENSG00000149925 | ALDOA | 15396.8 |
| ENSG00000182718 | ANXA2 | 13115.4 |
| ENSG00000039068 | CDH1 | 13110.1 |
| ENSG00000104267 | CA2 | 10551.6 |
| ENSG00000165376 | CLDN2 | 10228.7 |
| ENSG00000188488 | SERPINA5 | 10178.2 |

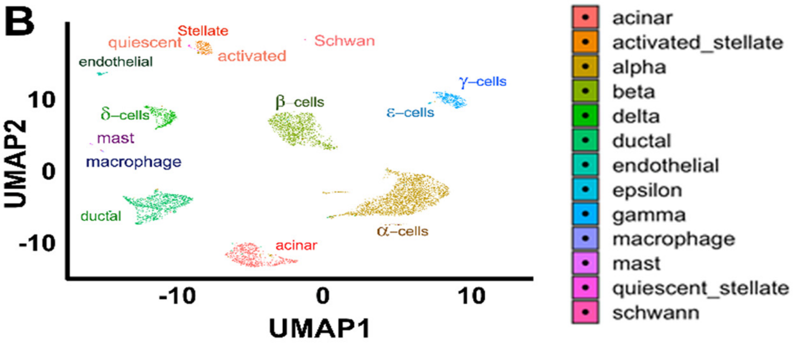


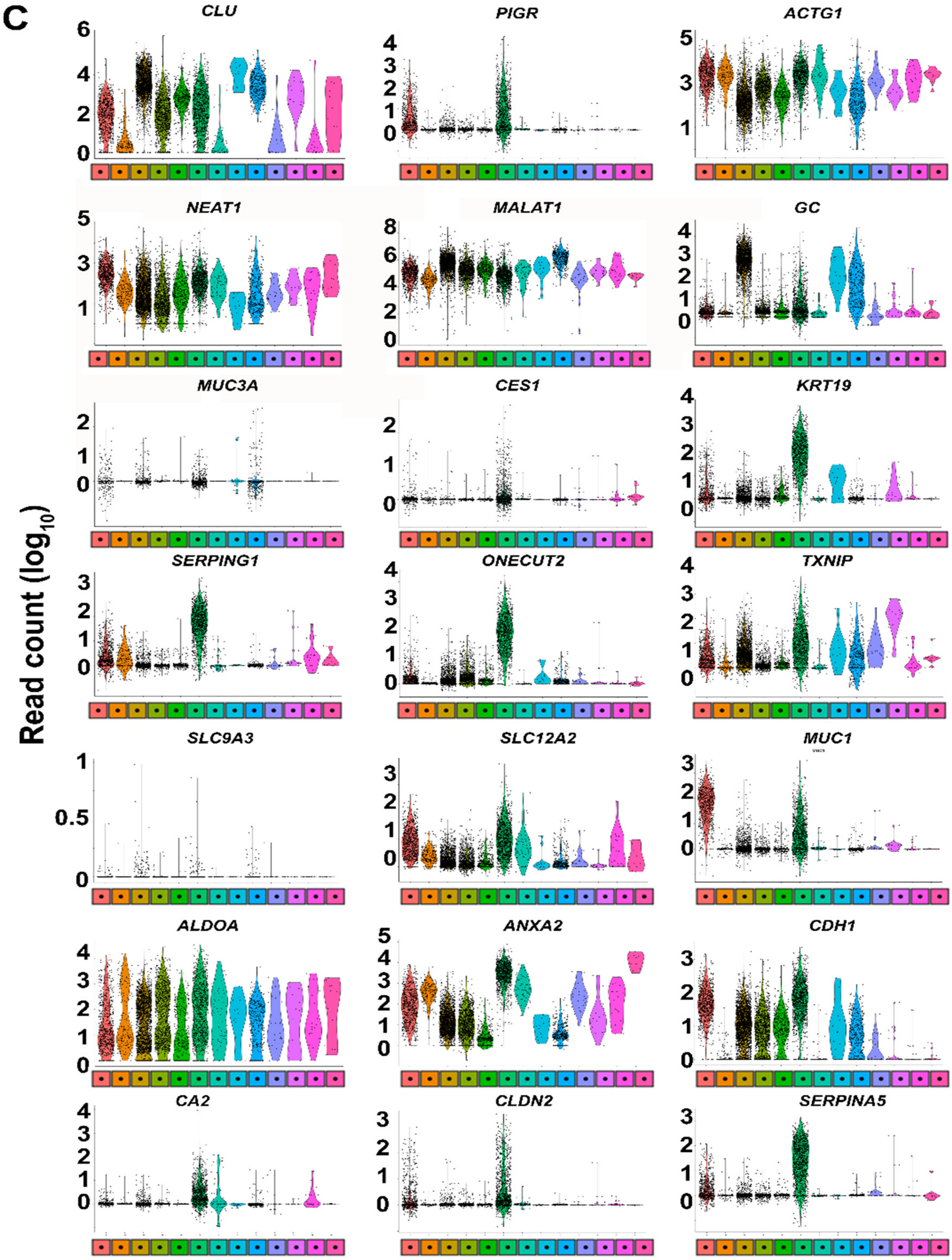

***<u>Fig. S4: Related to Fig. 3:</u>***

(**A**) Gene ID, gene symbol, and average read count of 21 highly abundantly human gallbladder-enriched gene transcripts. (**B**) UMAP: Uniform Manifold Approximation and Projection plot generated using Seurat (version 0.2.1) from human pancreatic single-cell RNA-sequencing (Panc8) data (see methods for details) is presented here. The single cells were clustered into 13 different pancreatic cell types, where each cluster is highlighted with a different colour. (**C**) Violin plots demonstrating the expression of each gene listed in panel A across all human pancreatic cell types presented in panel B. The Y-axis represents read counts (expression) of the gene transcript presented in each panel for the subset of human pancreatic cell type shown in panel B. Shape of the violin plot represents the density of individual data points and extend to min/max values. The colour of the violin plot reflects the colour and cell type from the UMAP plots in panel B.

50 **Fig. S5: Characterizing adult human gallbladder epithelial cells *in vitro***

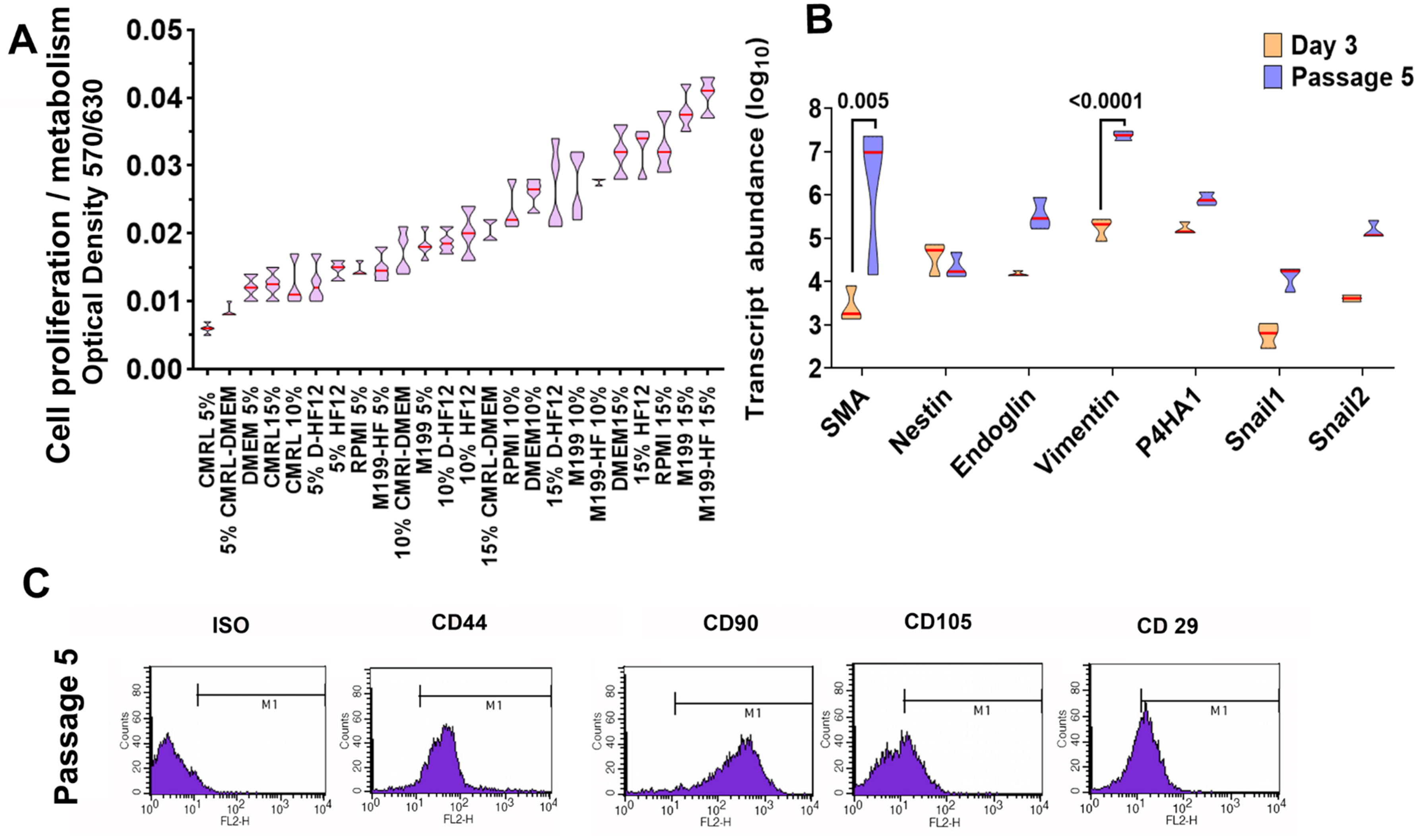


51

***Fig. S5: Related to Fig. 4:***

(**A**) Optical density/absorbance following an MTT assay in isolated gallbladder epithelial cells in the presence of different growth-supporting conditions is on Y-axis. Culture media and percentages (%) of fetal bovine serum (FBS) compositions are shown on the X-axis. Data are obtained from at least 3-4 different gallbladder preparations set up in a 96-well plate. (**B**) TaqMan®-based real-time quantitative (q)PCR for mesenchymal genes in freshly isolated gallbladder epithelial cells and *in vitro* propagated gallbladder-derived mesenchymal cells. Experiments are performed at least three times using different biological samples. Transcript abundance was calculated as described in methods and statistical significance between groups was assessed using two-way ANOVA with a p-value presented after adjustment using Sidak's multiple comparisons test. Polygons represent density distribution extending to min/max values with a horizontal red line at the median for Panel A and B. (**C**) Representative flow cytometry plots of gallbladder-derived mesenchymal cells at passage 5 for mesenchymal surface antigens. M1 gating indicates antigen-positive cells that are outside of the isotype control (ISO). The same acquisition and gating parameters are used while comparing isotype and antibody stained cells. Experiments were repeated at least three times with different gallbladder donor cell preparations.

### Supplementary Tables

**Table S1:** Gene assay IDs used for qPCR. List of TaqMan® primer/probe assays selected for real-time qPCR on the ViiA7 platform.

| Target species | Assay ID | Gene symbol | Gene name | Dye |
|---|---|---|---|---|
| Human | Hs02741908_m1 | *INS* | Insulin | FAM-MGB |
| Human | Hs00174967_m1 | *GCG* | Glucagon | FAM-MGB |
| Human | Hs00174949_m1 | *SST* | Somatostatin | FAM-MGB |
| Human | Hs01651425_s1 | *MAFA* | V-Maf Avian Musculoaponeurotic Fibrosarcoma Oncogene Homolog A | FAM-MGB |
| Human | Hs00360700_g1 | *NGN3* | Neurogenin 3 | FAM-MGB |
| Human | Hs00172878_m1 | *HES1* | Hairy and Enhancer of Split 1 (Hes Family BHLH Transcription Factor 1) | FAM-MGB |
| Human | Hs00236830_m1 | *PDX1* | Pancreatic And Duodenal Homeobox 1 | FAM-MGB |
| Human | Hs03003631_g1 | *18S* | Eukaryotic 18S rRNA | VIC-MGB |
| Mouse | Mm01259683_g1 | *Ins1* | Insulin-1 | FAM-MGB |
| Mouse | Mm00731595_gH | *Ins2* | Insulin-2 | FAM-MGB |
| Mouse | Mm00801712_m1 | *Gcg* | Glucagon | FAM-MGB |
| Mouse | Mm00436671_m1 | *Sst* | Somatostatin | FAM-MGB |
| Mouse | Mm00437606_s1 | *Neurog3* | Neurogenin 3 | FAM-MGB |
| Mouse | Mm01342805_m1 | *Hes1* | Hairy and enhancer of split 1 | FAM-MGB |
| Mouse | Mm00435565_m1 | *Pdx1* | Pancreatic And Duodenal Homeobox 1 | FAM-MGB |
| Mouse | Mm00447452_m1 | *Hnf1b* | Hepatocyte nuclear factor 1-beta | FAM-MGB |
| Mouse | Mm00448840_m1 | *Sox9* | SRY-Box Transcription Factor 9 | FAM-MGB |
| Mouse | Mm00433964_m1 | *Hnf4a* | Hepatocyte Nuclear Factor 4-Alpha | FAM-MGB |
| Mouse | Mm00468656_m1 | *Hlx* | H2.0-like homeobox | FAM-MGB |
| Mouse | Mm03302249_g1 | *Gapdh* | Glyceraldehyde-3-Phosphate Dehydrogenase | FAM-MGB |

**Table S2:** List of the TaqMan® primer/probe gene expression assays for real-time qPCR on the TLDA™ platform. All assays use FAM-MGB dye at 20x stock concentration. Housekeeping gene (*18S*) is highlighted in grey.

| | Assay ID | Gene Symbol | Gene Name |
|---|---|---|---|
| 1 | Hs00169631_m1 | *INSR* | Insulin Receptor |
| 2 | Hs00218236_m1 | *FOXJ2* | Forkhead Box J2 |
| 3 | Hs00174949_m1 | *SST* | Somatostatin |
| 4 | Hs00607978_s1 | *CXCR4* | C-X-C Motif Chemokine Receptor 4 |
| 5 | Hs00426835_g1 | *ACTA2* | Actin Alpha 2, Smooth Muscle |
| 6 | Hs00183740_m1 | *DKK1* | Dickkopf WNT Signaling Pathway Inhibitor 1 |
| 7 | Hs00175619_m1 | *PCSK1* | Proprotein Convertase Subtilisin/Kexin Type 1 |
| 8 | Hs00707120_s1 | *NES* | Nestin |
| 9 | Hs00355773_m1 | *INS* | Insulin |
| 10 | Hs00174967_m1 | *GCG* | Glucagon |
| 11 | Hs00240792_m1 | *FGFR2* | Fibroblast Growth Factor Receptor 2 |
| 12 | Hs00274931_s1 | *GAL* | Galanin And GMAP Prepropeptide |
| 13 | Hs00174139_m1 | *CD44* | CD44 Molecule (Indian Blood Group) |
| 14 | Hs00171403_m1 | *GATA4* | GATA Binding Protein 4 |
| 15 | Hs00264887_s1 | *POU3F4* | POU Class 3 Homeobox 4 |
| 16 | Hs00240871_m1 | *PAX6* | Paired Box 6 |
| 17 | Hs00185584_m1 | *VIM* | Vimentin |
| 18 | Hs00165775_m1 | *GLUT2 (SLC2A2)* | Solute Carrier Family 2 Member 2 |
| 19 | Hs00153380_m1 | *CCND2* | Cyclin D2 |
| 20 | Hs00232018_m1 | *GATA6* | GATA Binding Protein 6 |
| 21 | Hs99999901_s1 | *18S* | Eukaryotic 18S rRNA |
| 22 | Hs00606262_g1 | *HDAC1* | Histone Deacetylase 1 |
| 23 | Hs00173014_m1 | *PAX4* | Paired box 4 |
| 24 | Hs00360700_g1 | *NEUROG3* | Neurogenin 3 |
| 25 | Hs00232355_m1 | *NKX6-1* | NK6 Homeobox 1 |
| 26 | Hs00232128_m1 | *HLXB9* | Motor Neuron And Pancreas Homeobox 1 |
| 27 | Hs00192380_m1 | *SERPINI1* | Serpin Family I Member 1 |
| 28 | Hs00170285_m1 | *ICAM5* | Intercellular Adhesion Molecule 5 |
| 29 | Hs00168575_m1 | *P4HA1* | Prolyl 4-Hydroxylase Subunit Alpha 1 |
| 30 | Hs00164438_m1 | *ENG* | Endoglin |
| 31 | Hs00159922_m1 | *PCSK2* | Proprotein Convertase Subtilisin/Kexin Type 2 |
| 32 | Hs00179829_m1 | *FGFR3* | Fibroblast Growth Factor Receptor 3 |
| 33 | Hs00210096_m1 | *KRT23* | Keratin 23 |

| | | | |
|---|---|---|---|
| 34 | Hs00247426_m1 | *DKK3* | Dickkopf WNT Signaling Pathway Inhibitor 3 |
| 35 | Hs00187320_m1 | *HDAC3* | Histone Deacetylase 3 |
| 36 | Hs00170423_m1 | *CDH1* | Cadherin 1 |
| 37 | Hs00606991_m1 | *MKI67* | Marker Of Proliferation Ki-67 |
| 38 | Hs00167155_m1 | *SERPINE1* | Serpin Family E Member 1 |
| 39 | Hs00159598_m1 | *NEUROD1* | Neuronal Differentiation 1 |
| 40 | Hs00231032_m1 | *HDAC2* | Histone Deacetylase 2 |
| 41 | Hs00169777_m1 | *PECAM1* | Platelet And Endothelial Cell Adhesion Molecule 1 |
| 42 | Hs99999905_m1 | *GAPDH* | Glyceraldehyde-3-Phosphate Dehydrogenase |
| 43 | Hs00234422_m1 | *MMP2* | Matrix Metallopeptidase 2 |
| 44 | Hs00157705_m1 | *GLP1R* | Glucagon Like Peptide 1 Receptor |
| 45 | Hs00169851_m1 | *NCAM1* | Neural Cell Adhesion Molecule 1 |
| 46 | Hs00158126_m1 | *ISL1* | ISL LIM Homeobox 1 |
| 47 | Hs00277220_m1 | *GCK* | Glucokinase |
| 48 | Hs00233790_m1 | *ITGAV* | Integrin Subunit Alpha V |

**Table S3:** List of the TaqMan® primer/probe gene expression assays for real-time qPCR on the OpenArray™ platform. All assays use FAM-MGB dye at 20x stock concentration. Housekeeping gene (*18S*) is highlighted in grey.

| | **Assay ID** | **Gene Symbol** | **Gene Name** |
|---|---|---|---|
| 1 | Hs00173014_m1 | *PAX4* | Paired box 4 |
| 2 | Hs00271378_s1 | *MAFB* | V-Maf Avian Musculoaponeurotic Fibrosarcoma Oncogene Homolog B |
| 3 | Hs00359592_m1 | *NOVA1* | Neuro-Oncological Ventral Antigen 1 |
| 4 | Hs00240858_m1 | *PAX2* | Paired Box 2 |
| 5 | Hs03003631_g1 | *18S* | Eukaryotic 18S rRNA |
| 6 | Hs00240871_m1 | *PAX6* | Paired Box 6 |
| 7 | Hs00268388_s1 | *SOX4* | SRY-Box Transcription Factor 4 |
| 8 | Hs01001343_g1 | *SOX9* | SRY-Box Transcription Factor 9 |
| 9 | Hs00603586_g1 | *PTF1A* | Pancreas Associated Transcription Factor 1a |
| 10 | Hs00413554_m1 | *HNF6 (ONECUT1)* | Hepatocyte Nuclear Factor 6 (One Cut Homeobox 1) |
| 11 | Hs01001602_m1 | *HNF1β (TCF2)* | Hepatocyte Nuclear Factor 1-Beta (Transcription Factor 2) |
| 12 | Hs00167041_m1 | *HNF1α (TCF1)* | Hepatocyte Nuclear Factor 1-Alpha (Transcription Factor 1) |
| 13 | Hs00230853_m1 | *HNF4α* | Hepatocyte Nuclear Factor 4-Alpha |
| 14 | Hs00541450_m1 | *GLIS3* | GLIS Family Zinc Finger 3 |
| 15 | Hs00232355_m1 | *NKX6.1* | NK6 Transcription Factor Related, Locus 1 |
| 16 | Hs00159616_m1 | *NKX2.2* | NK2 Transcription Factor Related, Locus 2 |
| 17 | Hs00896294_m1 | *PROX1* | Prospero-Related Homeobox 1 |
| 18 | Hs01367669_g1 | *HES3* | Hairy And Enhancer Of Split 3 (Hes Family BHLH Transcription Factor 3) |
| 19 | Hs01922995_s1 | *NEUROD1* | Neuronal Differentiation 1 |
| 20 | Hs00892681_m1 | *GLUT1 (SLC2A1)* | Solute Carrier Family 2 (Facilitated Glucose Transporter), Member 1 |
| 21 | Hs01096908_m1 | *GLUT2 (SLC2A2)* | Solute Carrier Family 2 (Facilitated Glucose Transporter), Member 2 |

| | | | |
|---|---|---|---|
| 22 | Hs01564555_m1 | *GCK* | Glucokinase |
| 23 | Hs00846499_s1 | *UCN3* | Urocortin 3 |
| 24 | Hs00158126_m1 | *ISL1* | ISL LIM Homeobox 1 |
| 25 | Hs00171403_m1 | *GATA4* | GATA Binding Protein 4 |
| 26 | Hs00232018_m1 | *GATA6* | GATA Binding Protein 6 |
| 27 | Hs00292465_m1 | *ARX* | Aristaless Related Homeobox |
| 28 | Hs01005963_m1 | *IGF2* | Insulin Like Growth Factor 2 |
| 29 | Hs00264887_s1 | *POU3F4* | POU Class 3 Homeobox 4 |
| 30 | Hs00703572_s1 | *BHLHA15 (MIST1)* | Basic Helix-Loop-Helix Family Member A15 |
| 31 | Hs00907365_m1 | *HB9 (MNX1)* | Homeobox HB9 (Motor Neuron And Pancreas Homeobox 1) |
| 32 | Hs00158750_m1 | *LMX1.2 (LMX1B)* | LIM Homeobox Transcription Factor 1 Beta |
| 33 | Hs00892663_m1 | *LMX1.1 (LMX1A)* | LIM Homeobox Transcription Factor 1 Alpha |
| 34 | Hs01078080_m1 | *CDX2* | Caudal Type Homeobox 2 |
| 35 | Hs00793699_g1 | *GSX1* | GS Homeobox 1 |
| 36 | Hs00370195_m1 | *GSX2* | GS Homeobox 2 |
| 37 | Hs01116195_m1 | *EXO1* | Exonuclease 1 |
| 38 | Hs00170171_m1 | *REG3A* | Regenerating Family Member 3 Alpha |
| 39 | Hs01551078_m1 | *TLR3* | Toll Like Receptor 3 |
| 40 | Hs00358111_g1 | *PPY* | Pancreatic Polypeptide |
| 41 | Hs00230829_m1 | *AIRE* | Autoimmune Regulator |
| 42 | Hs01074053_m1 | *GHRL* | Ghrelin And Obestatin Prepropeptide |
| 43 | Hs01026107_m1 | *PCSK1* | Proprotein Convertase Subtilisin/Kexin Type 1 |
| 44 | Hs00159922_m1 | *PCSK2* | Proprotein Convertase Subtilisin/Kexin Type 2 |
| 45 | Hs00232764_m1 | *HNF3β (FOXA2)* | Hepatocyte Nuclear Factor 3-Beta (Forkhead Box A2) |

**Table S4:** List of primer sets used for ChIP DNA qPCR. Sequences start from 5' to 3'. Abbreviation: promoter (pro). Gene loci for which probe sequence is not provided are performed using SYBR green chemistry. TaqMan qPCR was used for human INS-275 and INS+1318.

| **Species** | **Gene locus** | **Forward primer** | **Reverse primer** | **Probe** |
|---|---|---|---|---|
| Mouse | Pdx1 A-I | CCAGTATCAGGGAGGACTATCA | TACCCAGCCATTAGGCAAGA | |
| Mouse | Pdx1 A-III | ACCGTGTCACCAAGTCAACCC | AGAGCCACCTGTGCCCGTCAA | |
| Mouse | Pdx1 A-IV | CTCTTCCTGATTCCCTGAAGTC | ACTAAGAGTGCTCTGGGCTCTG | |
| Human | INS pro | GTGGAAAGTGGTTTAGGTGAGGGT | ACCTGCTTGATGGCCTCTTCTGAT | |
| Human | PDX1 pro | CACACAACGAATGCCAGAGTTTCG | ACTGATCTCAGAGGGAACCCACA | |
| Human | NEUROG3 pro | AAGAGAGGCAGTGAAACACCAGGA | AACTCTCGGTTCCTCAAAGAGCCT | |
| Human | HES1 pro | TCCTCCTCCCATTGGCTGAAAGT | TTGGTGATCAGTAGCGCTGTTCCA | |
| Human | INS -275 | TGTGAGCAGGGACAGGTCTG | TCCTCAGGACCAGCGGG | 6FAM-CCACCGGGCCCCTGGTTAAGACTCTA |
| Human | INS +1318 | CAGCTGGAGAACTACTGCAACTAGA | GCTGGTTCAAGGGCTTTATTCC | 6FAM-CCGCCTCCTGCACCGAGAGAGA |

**Table S5:** List of genes obtained from RNAseq data that are either unique or common to gallbladder or beta cells. Provided as separate Excel sheet.
